# The Alzheimer’s Disease Risk Genes MS4A4A And MS4A6A Cooperate to Negatively Regulate Trem2 and Microglia states

**DOI:** 10.1101/2024.11.23.625001

**Authors:** Dalya Rosner, Jessica Sun, Angie Yee, Chaitanya Wagh, Anna Rychkova, Rita Cacace, Alice Buonfiglioli, Muhammed Alwahagri, Phil Kong, Marina Roell, Wei-Hsien Ho, Belvin Gong, Heidi Denton, Giacomo Muscarnera, Tim Meese, Malak El-Khatib, Daniel Bermingham, Adiljan Ibrahim, Julia Kuhn, Xiaoting Wang, Daniel Gulbranson, Herve Rhinn, Zia Khan, Ananya Mitra, Tina Schwabe, Karpagam Srinivisan, Ilaria Tassi, Lotje DeWitte, Renzo Mancuso, Hua Long, Peter Heutink, Sara Kenkare-Mitra, Arnon Rosenthal

## Abstract

Genetic variations in *MS4A4A* and *MS4A6A* are linked to the regulation of cerebrospinal fluid soluble TREM2 (sTREM2) levels and are associated with Alzheimer’s disease (AD) risk and progression. Using CRISPR knockout and MS4A4A-degrading antibodies in primary human microglia, non-human primates (NHP), and a xenotransplantation model of amyloid pathology, we provide evidence that *MS4A4A* and *MS4A6A* are negative regulators of both the transmembrane and soluble TREM2 proteins. They also negatively regulate microglia proliferation, survival, metabolism, lysosomal function, energetics, phagocytosis, and disease-fighting states. Mechanistically, we find that MS4A4A exerts negative regulation by interacting with MS4A6A and protecting it from degradation. MS4A6A in turn forms a complex with and blocks the co-receptor DAP12, which is required for the stability, cell surface localization, and signaling of TREM2 and other receptors. Taken together, the data indicate that *MS4A4A* and *MS4A6A* are cooperating, post-transcriptional negative regulators of TREM2 and microglial function, and potential drug targets for AD.

## Introduction

Microglia play a crucial role in maintaining brain health by containing misfolded proteins, removing damaged synapses and myelin, regulating astrocytes, oligodendrocytes, and endothelial cells, and modulating neuronal function^1^. With aging, due to detrimental genetic mutations and disease burden, microglia become dysfunctional, unable to counteract emerging pathologies, and display maladaptive stress response that may exacerbate Alzheimer’s disease (AD) pathology^2, 3^. Highlighting the importance of microglia, over 50% of the approximately 75 risk genes for AD have been shown to regulate microglia survival, proliferation and function^4–12^. A notable such AD risk gene encodes the single transmembrane protein TREM2^7, 10, 11^.

TREM2 is a damage-sensing receptor expressed on tissue macrophages and microglia in the brain^13^. It is activated upon binding to injury ligands, including phospholipids released from disintegrating membranes^14^, apoptotic cell debris, aggregated amyloid-β (Aβ) that leaches from amyloid plaques^15^, and misfolded proteins such as TDP43^16^ that are found in the AD brain. Ligand binding induces signaling through the TREM2 co-receptors DAP10 and DAP12, activating the SYK, PI3K, AKT, GSK3β, and mTOR pathway, and facilitating microglia survival, proliferation, migration, mitochondrial and lysosomal function, phagocytosis, and the transition to a disease-fighting state^13, 17, 18^.

A soluble form of human TREM2 (sTREM2) is found in the cerebrospinal fluid (CSF) and bloodstream^19, 20^. sTREM2 is produced either by protease cleavage^21, 22^ or alternative splicing^23^. High levels of sTREM2 in human CSF are thought to primarily indicate higher levels of transmembrane TREM2 on microglia^24^ have emerged as a predictor of lower AD risk and delayed disease progression^25, 26^. Longitudinal studies have shown that higher baseline levels of sTREM2 correlate with slower accumulation of Aβ plaques and Tau aggregates, slower decline in episodic memory and global cognition, slower hippocampal and cortical volume loss, delayed age of onset, better survival, and slower progression from mild cognitive impairment to full AD^24, 26–30^. Mendelian randomization analyses further demonstrate that lower sTREM2 levels are causally linked to worsening sporadic AD pathology^31, 32^.

A strong regulator of sTREM2 is located at the multi-gene *MS4A* locus, which encodes a family of tetraspanin proteins^6, 30, 33–35^. This locus is also associated with modulation of AD risk, age of onset, and CSF phospho-Tau levels^30, 36^. Recently, fine mapping of the locus has identified functional variants in both *MS4A4A* and *MS4A6A* each independently impacting CSF sTREM2 levels and AD risk^36^. *MS4A4A* and *MS4A6A* are predominantly expressed in myeloid cells in the periphery and microglia in the brain^37–39^. Variants in the *MS4A* locus that increase AD risk are associated with reduced CSF sTREM2 levels while protective variants were shown to correlate with elevated levels of sTREM2 in the CSF^30^. Little is known about the function of the *MS4A* proteins, though they were shown to sense long-chain fatty acid, steroid, and heterocyclic compounds and to possibly function as olfactory receptors^40^, and were implicated as ion channels in the regulation of immune response^41^ as switches of cell cycle and differentiation in health and disease^42^. Overexpression of MS4A4A has been suggested to elevate TREM2 in in macrophages^30^. However, the mechanism by which MS4A4A and MS4A6A regulate TREM2, their role in myeloid cell function, and whether protection from human AD is related to a loss or gain of MS4A protein function remains unclear.

Here, we show that genetic or pharmacological ablation of MS4A4A and/or MS4A46A, lead to the posttranscriptional elevation of soluble and membrane TREM2, to the modulation of transcripts and proteins that direct proliferation, survival, metabolism, lysosomal function, phagocytosis and to the adoption of disease protective state in primary macrophages, iPSC microglia, primary human microglia, cynomolgus monkey microglia and/or human microglia from amyloid-bearing mice.

Mechanistically, our work revealed that MS4A4A and MS4A6A form a complex that protects MS4A6A from degradation. MS4A6A is then able to establish an inhibitory complex with TYROBP/DAP12, the co-receptor required for stabilization, membrane localization, and signaling of TREM2 and other receptors, thereby preventing TREM2 membrane localization and signaling and possibly the activities of other DAP12 dependent receptors in microglia.

Taken together, these data identify the AD risk genes *MS4A4A* and *MS4A6A* as cooperative negative regulators of TREM2 and microglial functionality and reveal a new integrated mechanism through which a cascade of at least four AD risk genes (*MS4A4A, MS4A6A, TREM2, TYROBP/DAP12*), interact. The data further suggest that MS4A4A and MS4A6A could be promising therapeutic targets for AD.

## Results

### MS4A4A and MS4A6A form a complex that negatively regulates TREM2

We began our studies by determining whether MS4A4A and/or MS4A6A regulates TREM2. For this, *MS4A4A* or *MS4A6A* were knocked out (KO) using CRISPR/Cas9 in primary human macrophages and the resulting levels of membrane and soluble TREM2 were measured. *MS4A6A* KO led to a loss of MS4A6A protein but not of MS4A4A protein. In contrast, *MS4A4A* knockout led to the ablation of both MS4A4A and MS4A6A proteins (Figure 1A). The levels of *MS4A6A* mRNA were unaffected by *MS4A4A* knockout (Supplemental Figure 1), indicating that MS4A4A regulates MS4A6A protein post-transcriptionally. *MS4A4A* and *MS4A6A* KOs increased the levels of membrane TREM2 levels by 2-fold and 3.5-fold and the levels of sTREM2 by 9-fold and 11-fold, respectively (Figure 1B, C), supporting the idea that each of these MS4A proteins can negatively regulate TREM2 post-translationally.

**Figure 1:**
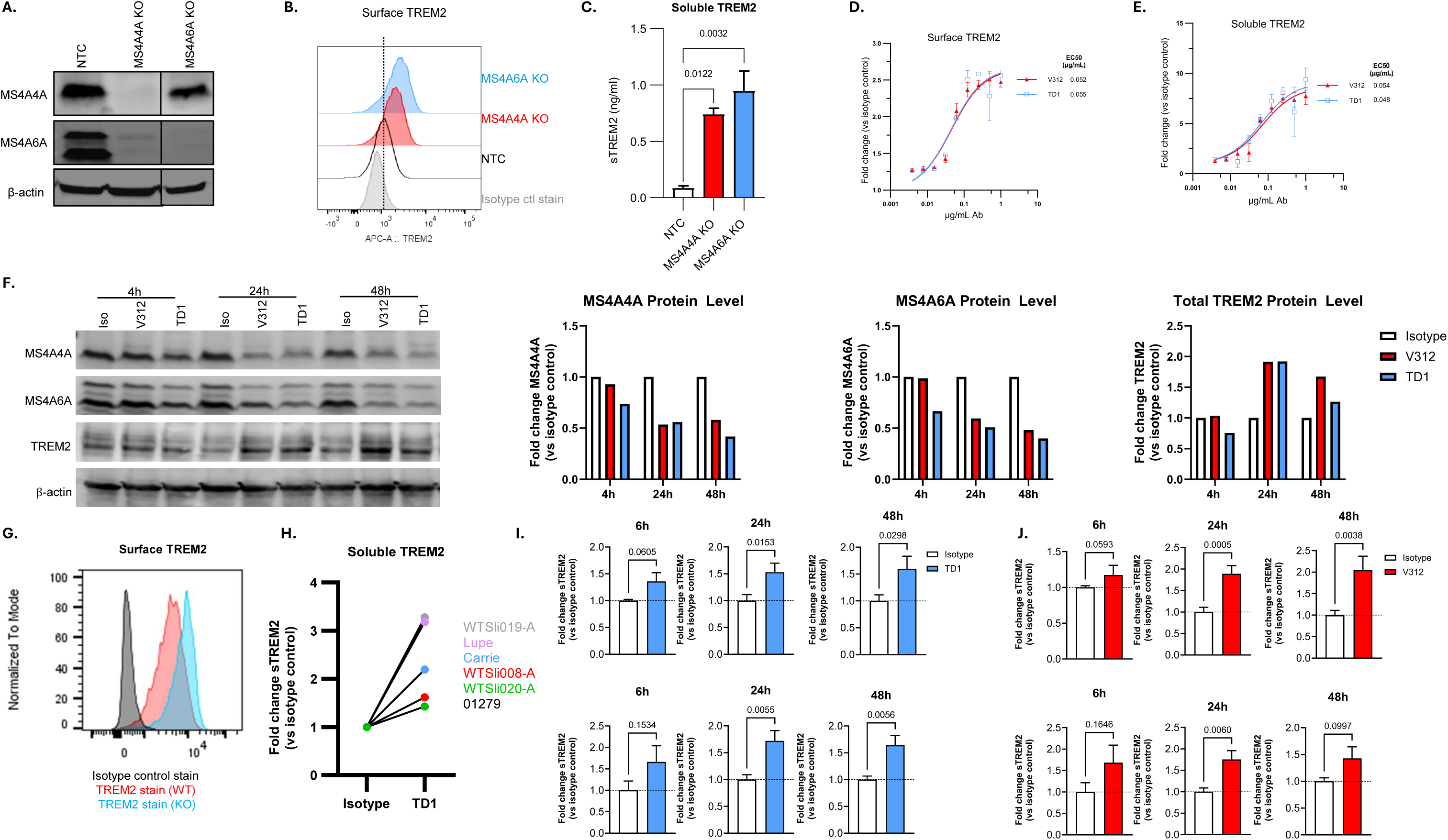
MS4A4A negatively regulates TREM2. (A) MS4A4A and MS4A6A were detected by Western from non-targeted control (NTC), MS4A4A KO and MS4A6A KO macrophages. Representative image of n=3 donors. (B) Surface and (C) soluble TREM2 were measured from NTC, MS4A4A KO and MS4A6A KO macrophages. Representative of n=4 donors. (D) Surface and (E) soluble TREM2 levels in macrophages (n=3 donors) treated with V312, TD1 or isotype control for 48 hours. (F) MS4A4A4, MS4A6A and TREM2 levels in macrophages treated with 1 ug/mL of V312, TD1 or isotype control antibodies for 4, 24 or 48 hours. Representative blot and representative quantification from n=4 donors. (G) Surface TREM2 in a WT or MS4A4A KO iMG line (Lupe). Representative of two experiments. (H) Soluble TREM2 from six iMG lines treated with 1 μg/mL TD1 or isotype control for 48 hours. (I, J) Solub;e TREM2 was measured from co-cultures of cerebral organoids and microglia from MSN38 and MSN9 lines treated with 1 µg/mL TD1 (I), V312 (J) or isotype control for 6, 24 or 48 hours. Data shows mean + SEM of at least 6 organoids per cell line. p values were calculated using T-tests.

To extend these findings, we developed two antibodies (TD1 and V312) that specifically recognize the cell surface human and NHP MS4A4A protein. We did not generate equivalent antibodies against MS4A6A because it is mainly intracellular^30, 43^ (Supplemental Figure 2). Macrophages treated with these antibodies showed an increase in surface TREM2 (>2-fold) and sTREM2 (>5-fold) within 48 hours of treatment, with an EC_50_ of ∼0.05 μg/mL (Figures 1D, E). Both antibodies downregulated MS4A4A and MS4A6A (Figure 1F) by ∼40% (V312) to ∼60% (TD1) compared to isotype-treated controls. Both antibodies achieved maximal downregulation of the MS4A4A protein and maximal elevated levels of TREM2 within 48 hours albeit with different kinetics.

Since microglia are the main cell type that express TREM2 and sTREM2 in the brain, we further investigated whether MS4A4A regulates TREM2 also in microglia. When *MS4A4A* was knocked out using CRISPR/Cas9 in several iPSC-derived microglia (iMG) lines, TREM2 levels increased ∼2.4-fold on the cell membrane (Figure 1G), and sTREM2 was elevated between 1.2- to 4.4-fold in the conditioned media of the different iMG lines (Figure 1H). Treatment of iMG co-cultured with neurons in 3-D cerebral organoids with TD1 or V312 also led to a 1.5- to 2-fold elevation in sTREM2 (Figure 1I, J). Altogether, these data confirm that MS4A4A negatively regulates TREM2 in both human macrophages and microglia.

### MS4A6A Negatively Regulates TREM2 by Blocking Its DAP12 Co-Receptor

Next, we undertook to understand the mechanism by which MS4A4A and MS4A6A negatively regulate TREM2. Previous studies using yeast-two-hybrid and FLIM-FRET methodologies suggested that MS4A4A and MS4A6A form a physical complex^39^. We extended these findings by showing that endogenous physiological MS4A4A co-immunoprecipitated with MS4A6A in primary human macrophages (Figure 2A), suggesting that they physically interact and mechanistically cooperate.

**Figure 2:**
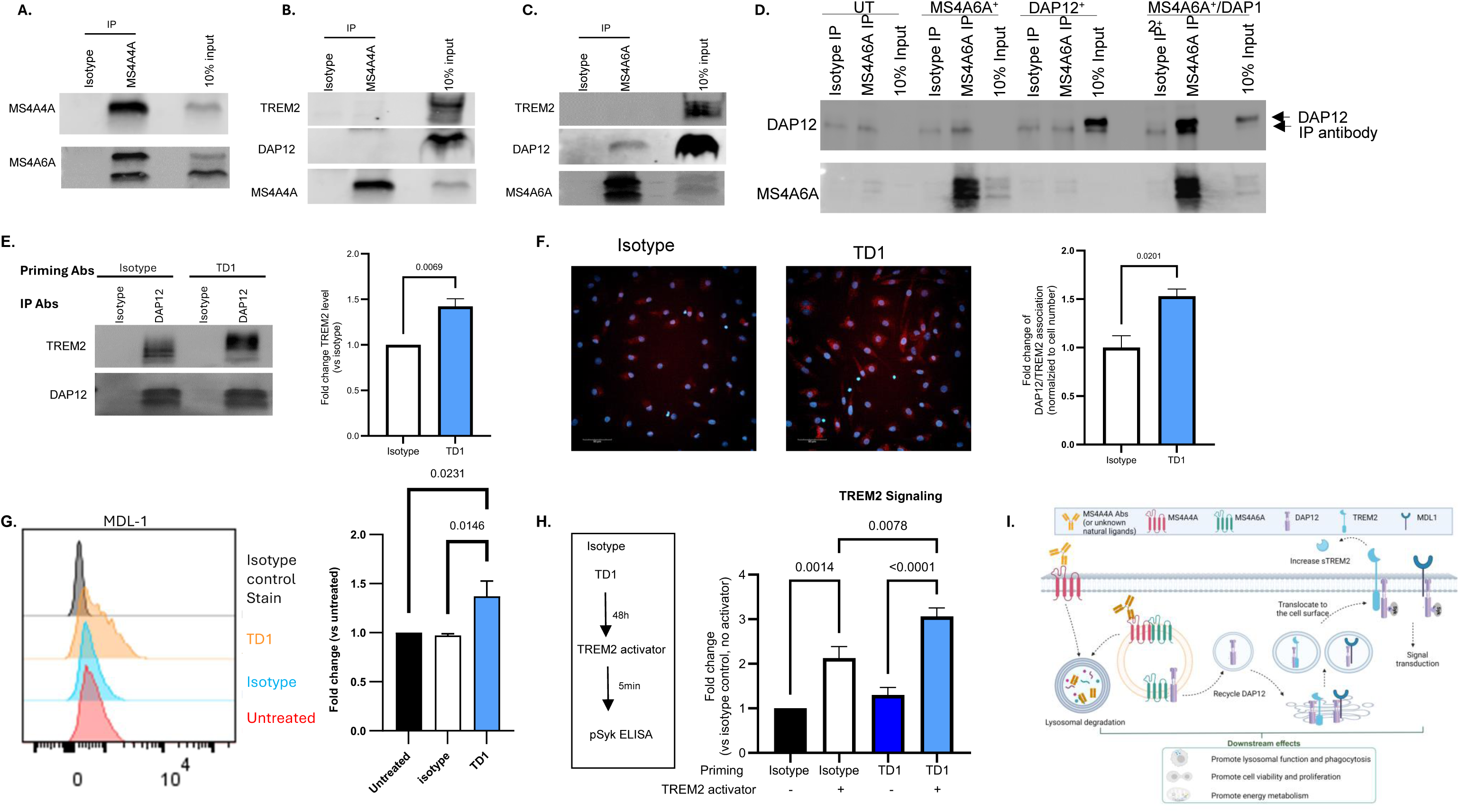
MS4A4A regulates TREM2 and downstream signaling through modulation of an MS4A6A/DAP12 axis. (A) MS4A6A was immunoblotted after MS4A4A immunoprecipitation. Representative image of n=3 macrophage donors. (B,C) TREM2 and DAP12 were immunoblotted from MS4A4A (B) or MS4A6A (C) immunoprecipitates. Representative image of n=3 donors. (D) Untransfected (UT) HEK293 and cells transfected with DAP12-GFP, MS4A6A-HA, or both were immunoblotted for DAP12 and MS4A6A after MS4A6A immunopreciptation. Whole cell lysate (10% immunoprecipitation input) was loaded as a control. Representative of n=2 experiments. (E) Macrophages were treated with 1 ug/mL TD1 or isotype control for 4 hours, then TREM2 and DAP12 were immunoblotted from DAP12 immunoprecipitates. n=3 donors. (F). Proximity ligation assay detecting the association between DAP12 and TREM2 after 48-hour treatment with 1 µg/mL TD1 or isotype control.. (G) MDL1 levels were measured on FACS after treatment with 1 µg/mL TD1 or isotype control for 48 hour. n=6 donors. (H) Macrophages were primed with 1 µg/mL TD1 or isotype control for 48 hours, then cells were treated with a TREM2 activator or a control for 5 minutes. Phosphorylated Syk was measured from the lysates. n=6 donors (I) Model of MS4A4A and MS4A4A-degrading antibodies mechanism of action on DAP12-associated receptors and effects of downstream functions. (D-G) Data in graphs show mean±SEM. p values were calculated by t-test (D,E) or one-way ANOVA (F, G).

We then tested the hypothesis that TREM2 may physically interact with either MS4A4A and/or MS4A6A. Surprisingly, TREM2 did not co-immunoprecipitate with either MS4A4A (Figure 2B) or MS4A6A (Figure 2C) even when weak interactions were chemically stabilized. Instead, immunoprecipitation of MS4A6A, but not of MS4A4A, led to co-immunoprecipitation of the TREM2 co-receptor DAP12 (encoded by *TYROBP)* (Figure 2B, C). To confirm the MS4A6A/DAP12 interaction, we transfected HEK293T cells with *DAP12* and *MS4A6A* constructs. When MS4A6A protein was immunoprecipitated from the doubly transfected cells, DAP12 co-immunoprecipitated. The interaction between MS4A6A and DAP12 was found to be independent of the presence of TREM2 or MS4A4A which are not expressed in HEK293T (Figure 2D). Since we found that MS4A4A stabilizes MS4A6A (Figure 1A, F), we undertook to test the hypothesis that MS4A4A may negatively regulate TREM2 indirectly, by increasing the availability of MS4A6A, which in turn forms a second, inhibitory complex with DAP12 and prevents DAP12 and TREM2 from interacting.

To this aim, we treated macrophages with TD1 for 4 hours to reduce the levels of MS4A4A and then quantified the amount of TREM2 that co-immunoprecipitated with DAP12 (Figure 2E). TD1-treated macrophages displayed 42% more TREM2 associated with DAP12 than isotype-treated controls. The TREM2 associated with DAP12 was at a higher molecular weight compared to free TREM2, consistent with the heavily glycosylated, mature form capable of trafficking to the cell surface that DAP12 has been shown to promote^44–46^. To further confirm that degradation of MS4A4A leads to the enhanced association between DAP12 and TREM2, we used a proximity ligation assay, which detects signals only when two proteins of interest are <40 nm apart (Figure 2F). When primary human macrophages that were treated with TD1 for 48 hours were stained for TREM2 and DAP12 in this assay, there was a 53% increase in TREM2 and DAP12 interaction. Degrading MS4A4A with TD1 also led to a 37% higher density of the pro-inflammatory, cytokine-regulating Myeloid DAP12-associating Lectin-1 (MDL-1) receptor on the macrophages cell surface (Figure 2G). These findings support the idea that degrading MS4A4A leads to subsequent degradation of MS4A6A, which in turn frees DAP12 to interact with TREM2 and other DAP12 dependent receptors on microglia.

We further measured whether the higher levels of TREM2 lead to an increase in Trem2 signaling. Macrophages were treated with the MS4A4A degrading antibody TD1 for 48 hours and were then stimulated with a TREM2-activating antibody for 5 minutes. Cells were then lysed, and the levels of phospho-Syk, the downstream signaling kinase of TREM2^17, 18^, were measured. Downregulation of MS4A4A led to a 45% increase in TREM2-dependent phospho-Syk (Figure 2H), supporting the hypothesis that MS4A4A are physiological negative regulators of both TREM2 cell surface levels and the resulting DAP12-dependent downstream signaling.

Taken together, these findings support a model whereby MS4A4A forms a complex with and stabilizes MS4A6A. This enables MS4A6A to form an inhibitory complex with DAP12 which prevents DAP12 from interacting with TREM2 and other DAP12 dependent receptors thereby restricting their stabilization, maturation, surface trafficking and signaling Figure 5I).

### MS4A4A is a Negative Regulator of Microglia Viability, Metabolism, Lysosomal Function, Phagocytosis, and Inflammatory States In Vitro

To test if MS4A4A impacts microglia states, we conducted functional assays on myeloid cells that were exposed to the MS4A4A degrading antibodies *in vitro*. Treatment of primary macrophages or iMG with TD1 or V312 led to a ∼2-fold increase in cell viability (EC_50_ TD1: 0.151±0.041 μg/mL; V312: 0.062±0.007 μg/mL) (Figure 3A, Supplemental Figure 4A), and a ∼3-fold increase in the production of the microglia survival and proliferation factor CSF1 (TD1 EC_50_ 0.149±0.084 μg/mL) (Figure 3B). TD1 and/or V312 treatment also elevated CSF1 levels in microglia isolated from post-mortem brains of non-demented individuals (TD1: 2-fold; V312: 2.5-fold) and AD patients (TD1: 3.5-fold; V312: 1.5-fold) (Figure 3C, Supplemental Figure 4B). Additionally, the MS4A4A-degrading antibodies also increased total ATP levels in macrophages by 30%, suggesting that the MS4A4A negatively regulates, survival, mitochondrial function and metabolic states in myeloid cells, either directly or through TREM2 (Figure 3D, Supplemental Figure 4C).

**Figure 3.**
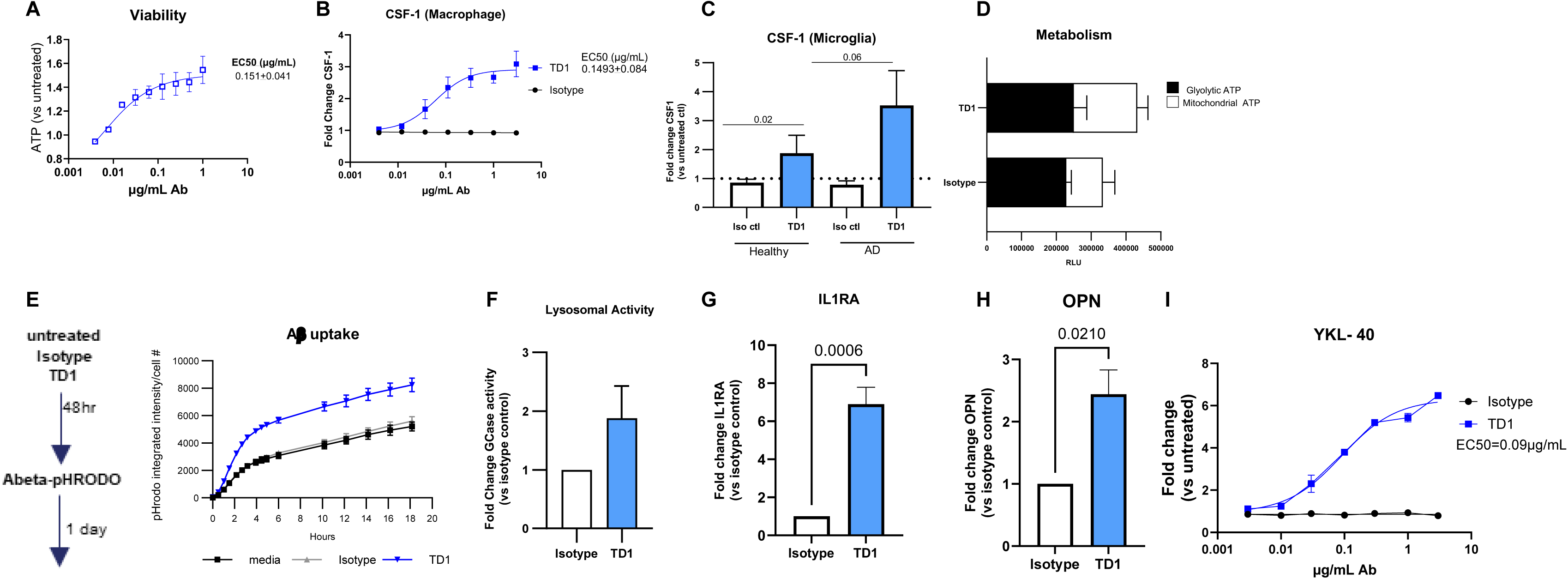
The MS4A4A-degrading antibody TD1 regulates viability, metabolism, lysosomal function, phagocytosis and cytokine production. (A) Macrophage viability (n=3 donors) and (B) CSF-1 production (n=4 donors) was measured after 48 hr treatment with TD1 or isotype control. (C) CSF-1 production was measured after 48 hr treatment with 10 μg/mL TD1 or isotype control from microglia isolated from postmortem brains of healthy controls (n=7) or AD patients (n=5). (D) Glycolysis and oxidative phosphorylation was measured in macrophages (n=2 donors) after 48 hr treatment with 1 μg/mL TD1 or isotype control. (E) Line 01279 iPSC microglia were treated with with 1 μg/mL TD1 or isotype control or left untreated for 48 hr, then uptake of pHRODO-labled Ab42 was measured. Representative of 2 independent experiments, (F) Lysosomal activity was measured from macrophages (n=3 donors) 48 hrs after treatment with 10 μg/mL TD1 or isotype control (G) IL1RA (n=4 donors) and (H) OPN (n=3 donors) were measured from macrophages after treatment for 48hrs with 1 μg/mL TD1 or isotype control; (I) YKL40 production was measured from macrophages after 48 hr treatment with TD1 or isotype control. Representative of n=3 donors. (A-I) Data shows mean±SEM. (A,B, I) EC50s were calculated with a 4PL non-linear regression. (C, G, H) p values were calculated with T tests. (with one-way ANOVA.

We also tested whether MS4A4A regulates microglia response to misfolded proteins. We found that iMG that were treated with either TD1 or V312 displayed a 63% increase in phagocytosed Aβ42 that was routed to the lysosome, as measured by increased signal from pH-sensitive pHRODO-labeled Aβ42 aggregates (Figure 3E). Treated iMG also displayed higher activity of the lysosomal enzyme glucocerebrosidase, which metabolizes the glycosphingolipid glucosylceramide into ceramide and glucose (TD1: 88%; V312: 43%) (Figure 3F, Supplemental Figure 4D) and whose deficiency constitute risk for multiple neurodegenerative disorders^47^. Taken together these findings support the idea that MS4A4A also regulates phagocytic and lysosomal activity in microglia.

We also tested whether MS4A4A regulates myeloid immune response. We found that inflammatory mediators such as IL1RA (Figure 3G, Supplemental Figure 4E), Osteopontin (OPN) (Figure 3H, Supplemental Figure 4F), and YKL-40 (Figure 3I) were elevated in macrophages treated with TD1 or V312. TD1 raised IL1RA by 6.9-fold, OPN by 2.4-fold, and YKL-40 by 6.5-fold, while V312 raised IL1RA by 4.7-fold and OPN by 2-fold. Altogether, these data suggest that MS4A4A negatively regulates inflammatory and immune response in myeloid cells.

### MS4A4A is a Negative Regulator of Microglia Viability, Metabolism, Lysosomal Function, Phagocytosis, and Inflammation in Non-Human Primates (NHP) In Vivo

To strengthen the hypothesis that MS4A4A is a physiological negative regulator of microglia states, we tested the effect of the MS4A4A-degrading antibody, TD1, in cynomolgus monkeys *in vivo*.

Animals received two doses of 80 mg/kg of TD1 separated by one month. Serum and CSF samples were taken throughout the study and brain samples were collected two days after the second dose (Figure 4A). TD1 accumulated in the serum and cerebrospinal fluid (CSF) throughout the study, indicating robust brain penetration (Figure 4B). Consistent with the in vitro findings, TD1 elevated sTREM2 over 2-fold in the serum and 3.5-fold in the CSF (Figure 4C). Total TREM2 levels were elevated 2-fold in extracts from the cortex and hippocampus (Figure 4D).

**Figure 4.**
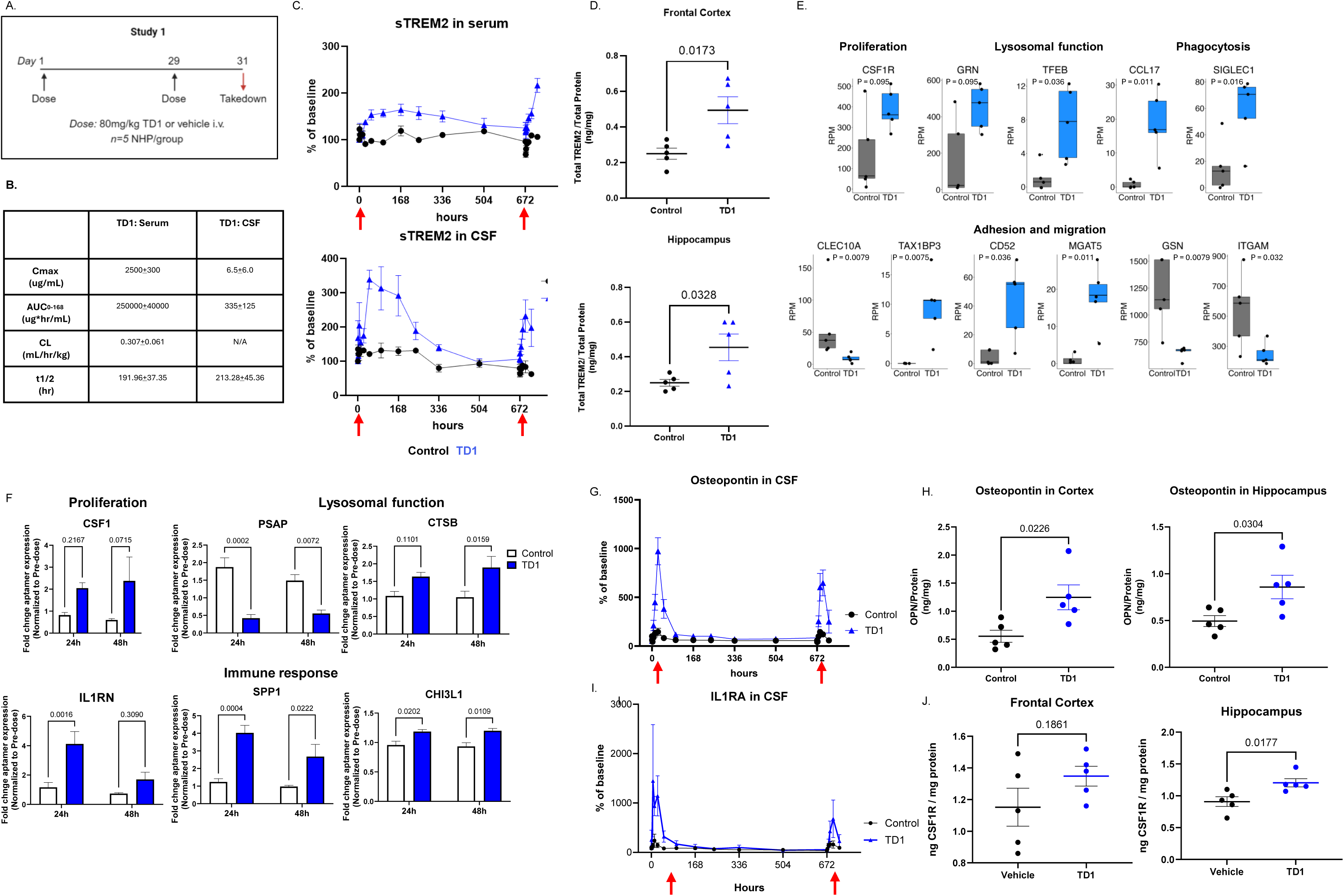
The MS4A4A- degrading antibody TD1 regulates TREM2 and microglial activation in non-human primates. (A) Study 1 design; (B) PK parameters (C) sTREM2 levels at the indicated time points in the serum (top) and CSF (bottom). (D) TREM2 protein levels in the frontal cortex and hippocampus 48 hours after the final dose of TD1. (E) Effect of TD1 on microglia gene expression in cortex at takedown. (F) Effect of TD1 on CSF proteins 24 and 48 hours after the first dose of antibody. (G) OPN and (I) IL1RA levels in the CSF at the indicated time points. (H) OPN and (J) sCSF1R protein levels in the frontal cortex and hippocampus 48 hours after final dose of TD1 Data shows mean±SEM for n=5 animals/group. p values were calculated with T-tests (D, H, JI) or Wilcoxon rank-sum test. (E) or two-Way ANOVA (F).

RNA-seq of isolated cortical microglia (Figure 4E) and proteomic analysis of CSF from TD1-treated cynomolgus monkeys (Figure 4F) demonstrated an effect of MS4A4A on microglia physiology that was in line with *in vitro* functional assays (Figure 3). Specifically, transcripts increased for regulators of proliferation (*CSF1R*), adhesion and migration (*CLEC10A, TAX1BP3, CD52, MGAT5, GSN, ITGAM*), phagocytosis (*CCL17, Siglec1*), and lysosomal function (*GRN, TFEB*) (Figure 4E). Changes to these transcripts and/or pathways correlated with alterations to CSF proteins, including up-regulation of proliferation signals (CSF1), lysosomal (PSAP, CTSB) and immune function (IL1RA, SPP1/OPN, CHI3L/YKL40) (Figure 4F). Quantitative ELISA-based assays revealed that Osteopontin/SPP1 was increased by 11-fold in the CSF (Figure 4G) and by 2-fold in extracts of the frontal cortex and hippocampus (Figure 4H). IL1RA levels were elevated by over 10-fold in the CSF (Figure 4I), while the soluble cleaved extracellular domain of CSF1R (sCSF1R) was increased by ∼1.3-fold in the hippocampal extracts (Figure 4J). In an independent study, animals were dosed 4 times over 50 days (Supplemental Figure 5A), TD1 also increased cortical microglia proliferation and total microglia numbers as measured by immunostaining for Ki67 (Supplemental Figure 5B, C).

To determine how well modulation of MS4A4A is tolerated, TD1 was administered to NHP at doses up to 257 mg/kg weekly for four weeks and multiple physiological outcomes were measured. TD1 was found to be generally safe, with no drug-related mortality or adverse findings related to clinical parameters, anatomy, ophthalmology, neurology, electrocardiograms (ECGs), blood pressure, body weight, food consumption, blood cytokine levels, or peripheral blood immunophenotyping. These findings support the conclusion that MS4A4A may have minimal effects on homeostatic physiology and that the observed effects of TD1 on the myeloid system are not a byproduct of a general stress response.

### Inhibition of MS4A4A Induces Protective Human Microglia in a Rodent AD Model

We next investigated the potential role of MS4A4A in an AD model. For this, 4-day-old *AppNL-G-F Rag2−/− Il2rγ−/− hCSF1KI* mice, which express pathological levels of amyloid-β (Aβ), were transplanted with human microglia following the MIGRATE protocol, as described previously^48, 49^. Six months later, when these mice exhibited extensive Aβ deposition, they were treated bi-weekly for three months with either V312 (80 mg/kg) or an isotype control (80 mg/kg) (Figure 5A). At nine months of age, human microglia were isolated by fluorescence-activated cell sorting (FACS), gating for CD11b+ and hCD45+ cells. Border-associated macrophages, low-quality cells, and doublets were removed, and single-cell RNA sequencing (scRNA-seq) was performed. The final dataset included 11,167 high-quality human microglia, which were classified into six transcriptional profiles, as described prebiously^48^: Homeostatic Microglia (HM), Cytokine Response Microglia (CRM), Interferon Response Microglia (IRM), Disease-Associated Microglia (DAM), Antigen-Presenting, HLA-expressing Microglia (HLA-M), and Ribosomal Microglia (RM) (Figure 5B, Supplemental Figure 6).

**Figure 5:**
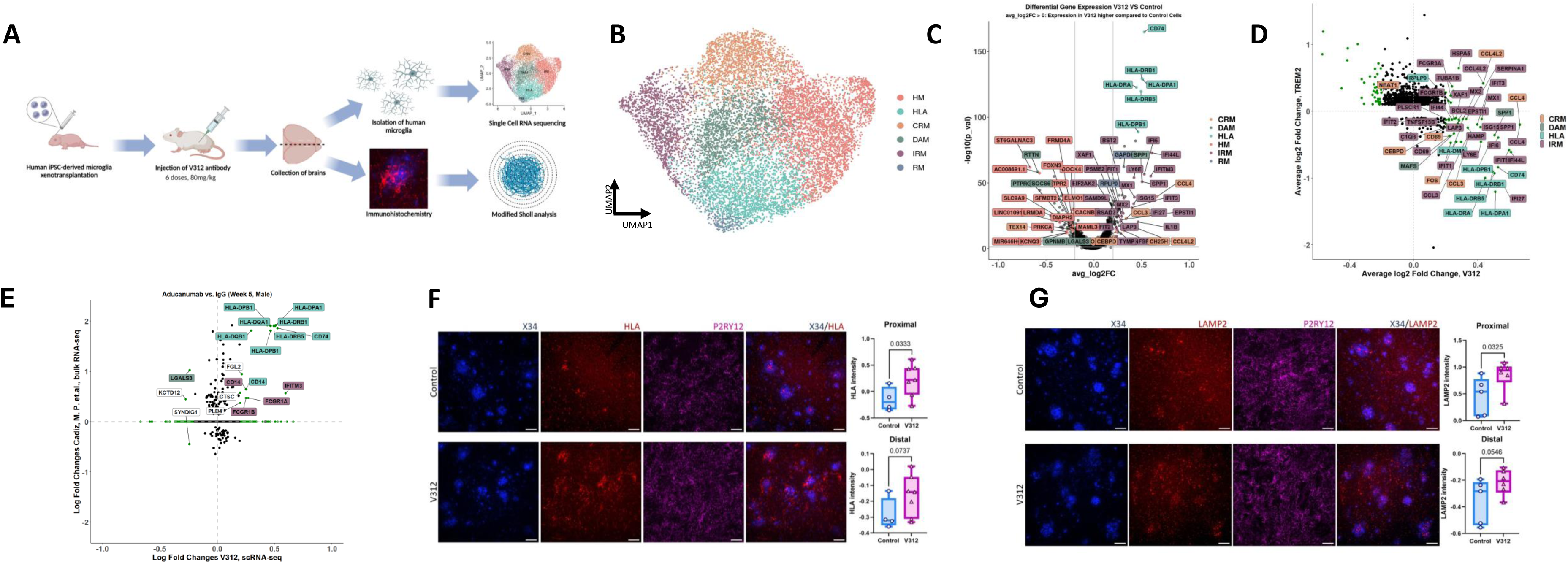
The MS4A4A-degrading antibody V312 induces microglia towards protective HLA+ cluster. (A) Experimental design of the study (V312 n=10 or isotype control n=8. (B) UMAP plot visualizing 11,167 high quality single human microglial cells sorted from mouse brain (CD11b^+^ hCD45^+^) after treatment with V312 or isotype control. (C) Volcano plot of differentially expressed genes average (log2Fold Change ≥ 0.2 and adjusted p-value ≤ 0.05) where each dot represents a cell. Labelled genes are differentially expressed and part of the top 30 marker genes of a cluster ranked on average log2Fold Change; (D) Comparison of V312 treatment with *TREM2^KO^* dataset^41^ (E) Comparison of our differentially expressed genes with selected genes from aducanumab-treated animals in ^42^ showing enhanced MHC (HLA) response male mice.(F,G) Representative images (scale bar 50 μm) stained for P2RY12 (HM microglia) and X-34 (Aβ plaques) and either HLA-DR/DQ (HLA microglia) (F) or LAMP2 (G) as proxy for phagocytosis. Quantification of HLA (F) and LAMP2 (G) markers at the site of Aβ plaques was performed with modified Sholl analysis. n=8 isotype control-treated mice and n=10 V312-treated mice. p values were calculated with one-tailed t-test.

Trajectory analysis previously showed that, in the presence of Aβ deposition, HM transition into DAM, which can further transition into HLA-M and IRM microglia subsets^48^. The HLA-M subset, characterized by the expression of MHC genes as well as *MS4A6A*, is unique to human microglia that were exposed to Aβ^48^ and to microglia from AD models treated with the anti-Aβ antibody aducanumab^50^, which clears Aβ plaques by recruiting microglia. The HLA-M subset is therefore thought to be protective. We found that V312 treatment upregulated several genes typical of the HLA-M cluster, including *CD74*, *HLA-DRA*, *HLA-DRB1*, *HLA-DRB5*, *HLA-DPA1*, and *HLA-DPB1* (Figure 5C, Supplemental Excel file). These findings support the idea that pharmacological degradation of MS4A4A in vivo leads to an increase in the prevalence of HLA-M protective microglia in response to Aβ pathology and are consistent with the findings that treatment of iMG with either TD1 or V312 led to a 63% increase in phagocytosed Aβ42 *in vitro* (Figure 3E).

DAM and HLA-M microglia subsets were shown to be dependent on TREM2: They fail to develop in microglia that carry the *TREM^R47H^* loss-of-function AD risk variant, or in TREM2*-*deficient microglia^48^. Consistent with these findings and the hypothesis that MS4A4A is a negative regulator of TREM2-dependent protective microglia, we observed that V312 boosted transcripts that were suppressed in *TREM2* KO microglia (Figure 5D).

When comparing transcriptomes of V312-treated and aducanumab-treated animals, we observed similar changes (Figure 5E, Supplemental Figure 7) suggesting that anti-Aβ therapeutics and MS4A4A-degrading antibodies induce a similar class of protective microglia. Immunostaining of brain tissue from *AppNL-G-F Rag2−/− Il2rγ−/− hCSF1KI* mice confirmed that V312 enriched for HLA-positive microglia (HLA-M-DQ/DR), particularly in proximity to Aβ plaques (Figure 5F, Supplemental Figure 8A). Immunostaining for lysosome-associated membrane protein-2 (LAMP2), a marker of phagocytic and endo-lysosomal activity, further showed a V312-dependent increase in microglial LAMP2 signal near Aβ plaques (Figure 5G, Supplemental Figure 8B), suggesting that the induced HLA-M microglia subset is associated with increased phagocytic and lysosomal activity.

Consistent with the findings that TREM2 signaling, although necessary, is not sufficient to induce all terminal microglial types^51^ and with prior reports that DAM phenotype requires two activation steps^52^, V312 did not alter the DAM microglia cluster at any distance from the plaques, as shown by IHC staining with hCD9, a known DAM signature marker (Supplemental Figure 9). Together, these findings support the hypothesis that MS4A4A negatively regulates microglial response to amyloid pathology.

## Discussion

*TREM2* itself is now recognized as a major risk gene for AD and a critical immune-stimulatory receptor for microglia with over 40 risk variants have been identified (https://www.alzforum.org/mutations/trem2). The *MS4A* locus has been linked either directly or through *TREM2* to both risk and progression of AD and appear to be a potent regulators of cerebrospinal fluid (CSF) sTREM2 levels^6, 30, 33–35^. Using CRISPR/CAS9 based genetic ablation and MS4A4A-degrading antibodies, in primary human macrophages, human iPSC microglia, human primary microglia, *in vitro* and NHP as well as a rodent AD model carrying human microglia *in vivo,* we provide evidence that MS4A4A and MS4A6A negatively regulate TREM2 and likely additional DAP12 dependent signaling receptors in microglia. At least in part through negative regulation of DAP12 dependent receptors such as Trem2, MS4A4A and MS4A6A modulate aspects of microglia survival, proliferation, lysosomal activity, energetics and state changes associated with the response to AD pathology. The gene expression signatures and microglia states that MS4A4A/ MS4A6A negatively regulate and those that TREM2 and anti Aβ antibodies positively regulate overlap, supporting the idea that these pathways intersect in modulating protective microglia response to AD.

Mechanistically, we found that MS4A4A is in a physical complex with MS4A6A and prevents its degradation. MS4A6A in turn forms an inhibitory complex with DAP12, a single transmembrane receptor which is essential for signaling, stabilization, cell membrane localization and intracellular trafficking of TREM2 and other receptors on microglia^53–56^. Physiologically, MS4A4A and MS4A6A may act as immune checkpoints to prevent premature microglial activation in response to low levels of damage-associated lipid ligands that may be generated during normal brain activity. In AD, MS4A4A and MS4A6A may exert excessive innate immune check point activity and prevent microglia from effectively responding to emerging disease pathologies.

Our work revealed a signaling pathway encompassing four risk genes for AD, *MS4A4A, MS4A6A, TREM2* and *TYROBP/DAP12* as well as a novel post-translational regulatory mechanism. MS4A4A and MS4A6A appear to negatively regulate microglia functions upstream of TREM2 and other DAP12 dependent receptors placing them as key negative modulators of myeloid cell function. As microglia immune check points, MS4A4A and MS4A6A are promising therapeutic targets for AD.

## Materials and Methods

### Primary human macrophage differentiation and culture

 Whole blood was incubated with RosetteSep Human Monocyte Enrichment Cocktail (StemCell Tech Catalog #15068) for 30 minutes, then mixed with Ficoll-Paque Plus (Cytiva #171404003) and monocytes were isolated using Sepmate tubes (StemCell Technologies #85460). After any residual RBCs were lysed with ACK, cells were plated at 10^6^/mL in macrophage media (RPMI + 10% FBS (Fisher # SH3007103HI), 1% HEPES (Fisher #15630080), 1% Penicillin/Streptomycin (Fisher #15140122), 1% sodium pyruvate (Fisher #15140122), 1% MEM NEAA (Fisher #15140122), 1% Glutamax (Fisher # 35050061) supplemented with 100 ng/mL of M-CSF (Biolegend #574808). After six days, cells had differentiated to macrophages and were re-plated for subsequent experiments. To re-plate, media was removed, cells were incubated for five minutes with PBS + 3mM EDTA, and then lifted. Cells were cultured in fresh macrophage media without addition of M-CSF at a density of 10^6^ cells/mL.

### iPSC derived microglia for 2D culture

Line 01279 (Fujifilm) were purchased pre-differentiated to microglia. Lines WTSli008-A, WTSli020-A and WTSli019-A (EBSC), Carrie (Thermo-Fisher), Lupe (Reprocell), and Pgp1 (Synthego) were differentiated in-house according to standard protocols^52^. In brief, to form embryoid bodies 1.5X 10^6^ iPSC cells were cultured in EB microglia media (Stem Cell Base Media (Stem Cell Technology #100-1130) +100ng/mL VEGF (Proteintech #HZ-1038)+40ng/mL SCF (Peprotech #300-07)+100ng/mL BMP4 (Peprotech #120-05)) + 10 mM Rock-i (only for the first 24 hours) (Peprotech #1293823) on Aggrewell 800 plates (Stem Cell Technology #34811) that had been pre-treated with anti-adherent solution (Stem Cell Technology #07010). After feeding cells daily for three days, embryoid bodies were transferred to a T150 flask pre-coated with 0.5% w/v autoclaved gelatin water solution and cultured in microglia factory solution (XVIVO (Lonza #02-053Q) + 25g/mL IL-3 (Peprotech #200-03)+100ng/mL M-CSF (Peprotech #300-25)+2Xb-mercaptoethanol (Gibco #21985023)+1X Glutamax (Gibco #35050061)) in order to induce microglia precursors. Flasks were fed on days 7, 14, 21, 24, 28, and 32 after feeding. On day 35, cells were harvested for differentiation, re-suspended at 1.44X 106 cells/mL in maturation media (Advanced DMEM/F12 (Gibco #12634010) +0.5X N2 (Gibco #17502048)+1X Glutamax+100ng/mL IL-34 (Proteintech #HZ1316)+10ng/mL GM-CSF (Peprotech #300-03)+2X b-mercaptoethanol+25ng/mL TGFb (Peprotech #100-21)+ 1.5 μg/ml cholesterol (Sigma #C3045), 1 ng/ml gondoic acid (Sigma #E3635), 100 ng/ml oleic acid (Sigma #O1008)) and seeded onto plates pre-coated with 1% Geltrex (Gibco #A14133-02). Cells were fed three times/week for two weeks and then were ready for further assay. CRISPR-Cas9 mediated KO of MS4A4A used guide RNA sequences GAAUGGAACAGGCCAUGCCA, and KO was validated by sequencing with GCCTATTCCACTTCCCCAGC primers.

### Organoid cultures

Cortical brain organoids were cultured using the STEMdiff^TM^ Cerebral Organoid Kit (#08570, #08571). The MSN (Mount Sinai Normal) −38 and −9 lines were obtained from the Stem Cell Engineering Core Facility at Icahn School of Medicine at Mount Sinai. At mature age, organoids were co-cultured with microglial progenitors to obtain microglia-containing cerebral organoids, as detailed in^57^. Co-cultures were stimulated with 5 µg/mL of antibody, and sTREM2 was measured by MSD at 6-, 24- and 48-hour time points.

### Antibody generation

All immunizations were performed at Alector in compliance with Institutional Animal Care and Use Committee (IACUC) and OLAW guidelines and regulations. Up to ten BALB/c or SJL mice at the age of 7–9 weeks were immunized using either DNA or a combination of cell and DNA against MS4A4A or MS4A6A. DNA immunizations were performed via hydrodynamic tail vein injections. Lymphocytes were isolated from the immunized animals and fused with either P3X63Ag8.653 (CRL-1580, American Type Culture Collection) or SP2/mIL6 (CRL-2016, American Type Culture Collection) mouse myeloma cells via electrofusion, selected in HAT media, and single colonies were isolated using the Clonepix2 system (Molecular Devices). After further expansion, supernatants were screened for binding by FACS, and positive clones were sequenced.

### Apparent KD measurement

20,000 U937-MS4A4A cells were blocked in FACS buffer (PBS+1% BSA) + 33% mouse serum, then incubated for 3 hours on ice with 200 mL of anti-MS4A4A antibodies with pipetting every 30 minutes. After washing, cells were incubated for 15 minutes on ice with 100 mL of 1:500 APC-F(ab’)2 fragment goat anti-human IgG (Jackson Immunoresearch #109-136-098) diluted in FACS buffer. Cells were then washed, fixed for 20 minutes with 4% paraformaldehyde, then washed again and run through a Canto flow cytometer (BD). Mean fluorescence intensity values measured by the flow cytometer were analyzed on Prism (GraphPad) or similar statistical programs, using the following equation (or similar 4-parameter fit) described in ^58^:

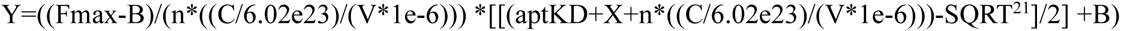

Where the following values are constrained according to experimental conditions: C = number of cells per well (20,000)

V = total volume of staining in microliters

n = number of receptors on cell surface, estimated to be 100,000 here

### Co-immunoprecipitation (Co-IP)

Cells were treated with chemical cross-linker, 1 mM dithiobis[succinimidylpropionate] (DSP, Thermo Fisher # A35393) on ice for 1h and stopped the reaction with quench buffer (1M Tris, PH 7.5, Thermo Fisher #15567027) for 5 min incubation at room temperature. Cells were washed with PBS three times before lysis with IP-lysis buffer (50 mM Tris HCL, 150 mM NaCl, 1 mM EDTA, 1.5 mM MgCl_2_, 10% Glycerol, 1% Triton X-100, 50 mM NaF, 1% n-Dodecyl β-D-maltoside (Sigma #D641) Halt protease and phosphatase inhibitor cocktail (Thermo-Fisher #1861281), PH 8) at a density of 10^7^ cells/500 mL lysis buffer. After preclearing with Protein G Sepharose (Sigma-Aldrich # P3296), cell lysates were incubated with Protein G Sepharose coated with 3 µg of antibodies against MS4A4A (Alector, 4A-Ab3), MS4A6A (Alector, 3G08), DAP12 (Santa Cruz Biotechnology #sc-133174) or TREM2 (R&D Systems #MAB17291) with gentle rotation at 4°C overnight per the manufacturer’s instructions. Immunoprecipitation isotype control samples were prepared in parallel with species-matched IgG. Following the incubation, immunoprecipitation samples were washed five times with 1mL of NET buffer (50 mM Tris-HCL, 150 mM NaCl, 1 mM EDTA, 1% Triton X-100, PH 7.5). The immunoprecipitated protein was solubilized with the elution buffer (1:1 mixture of NET buffer with 4X Laemmli buffer (Bio-Rad #1610747), and 10% β-mercaptoethanol, Sigma-Aldrich #444203). Meanwhile, 10% of the cell lysate or flowthrough of immunoprecipitation isotype control was used as an input control.

### Western Blot

Whole cell lysates were prepared with IP-lysis buffer and mixed with Laemmli buffer and 10% 2-mercaptoethanol without boiling. Whole cell lysates or immunoprecipitation samples were separated using Any kDa Mini-PROTEAN® TGX Stain-Free™ gels (Bio-Rad # 4568123 and #4568125) and transferred to PVDF membrane using the Trans-Blot Turbo kit (Bio-Rad #1704156).The membrane was blocked with 5% milk in 1X TBS with 0.1% Tween for 1 hour at room temperature and then incubated with primary antibodies against MS4A4A (1:300 dilution, Alector, Clone Gigi), MS4A6A (1:1,000 dilution, Abcam #ab189983), DAP12 (1:1,000 dilution, Santa Cruz Biotechnology #sc-133174 or Cell signaling Technology #12492) or TREM2 (1:1,000 dilution, R&D Systems #BAF1828) overnight at 4°C. Membranes were exposed to the appropriate horseradish peroxidase–conjugated secondary antibodies, followed by detection with SuperSignal West Femto (Thermo-Fisher #34096) or SuperSignal West Pico Plus (Thermo-Fisher #34580) chemiluminescent substrates. For whole cell lysates, equal protein loading was controlled by re-probing the membrane with anti–β-actin-HRP antibody (1:1,000 dilution; Santa Cruz Biotechnology #sc-47778-HRP). Chemiluminescence was detected using the ChemiDoc MP Imaging System (Bio-Rad) and Image J (NIH) was used for Western blot signal quantification.

### FACS

10^5^ cells were stained in duplicate or triplicates. Cells were washed in FACS buffer (PBS + 5% FBS + 0.1% sodium azide), then stained in FACS buffer + 1:50 Human TruStainX (Fc receptor block (Biolegend #422302) with antibody or isotype control for 30 minutes at 4C. Cells were then washed twice and re-suspended in FACS buffer + 1:2000 DAPI (Biolegend #422801). Cells were run through a Canto or Fortessa flow cytometer (BD). FlowJo software (Treestar) was used to calculate the MFI from a live (DAPI^−^), single-cell gate. Directly conjugated antibodies used were MDL-1-PE (Clone 14H4-1-2; 1:200; Biolegend # 371705), mouse IgG2b-PE isotype control (Clone MPC-11; 1:200; Biolegend #400312), TREM2-DL650 (Clone T22; 1:200 Alector), and Synagis mouse IgG1-DL650 (1:200; Alector). TREM2 and its isotype control antibody were labeled with DyLight-650 using a Microscale Antibody Labeling kit (Thermo-Fisher #84536). Indirectly conjugated antibodies used were V312-mIgG1 (Alector), TD1-mIgG1 (Alector), Synagis-mIgG1 (Alector) followed by 1:500 APC-F(ab’)2 goat-anti-mouse IgG (Jackson Immunoresearch #115-136-071.

### Transfections

HEK293 cells were transfected with 100 mg of DNA using the Lipofectamine 3000 transfection reagent (Thermo-Fisher # L3000015) and cells were lysed 24 hours later. Constructs were designed in Vector Builder with pCDNA.3 plasmids and the constructs generated were MS4A4A-isoform 1-HA, MS4A6A-isoform 2-HA, and DAP12-GFP. The HA construct was based on the “Snorkel” tag described in ^54^.

### Determination of antibody cross-reactivity

To confirm that MS4A4A antibodies did not bind to MS4A6A and vice versa, HEK293 cells were transfected with either MS4A4A-HA or MS4A6A-HA or mock transfected. To test specificity of Western antibodies, HA was IP’ed, then immunoblots were performed with antibodies used for MS4A4A (Alector; Clone Gigi), MS4A6A (Abcam #ab189983) or HA Western. MS4A4A antibody only detected protein in the MS4A4A-transfected cells and MS4A6A antibody only detected protein in the MS4A6A-transfected cells (Supplemental Figure 3C). To test specificity of IP antibodies, IPs were performed with MS4A4A (Alector; Clone 4A-Ab3) and MS4A6A (Alector; Clone 3G08) antibodies, then immunoblots were performed with antibodies used for MS4A4A (Alector; Clone Gigi), MS4A6A (Abcam #ab189983) or HA Western. Transfected protein (detected by HA) was only detected in MS4A4A-transfected cells that had an MS4A4A IP or in MS4A6A-transfected cells that had an MS4A6A IP (Supplemental Figure 3D). To test specificity of functional antibodies, FACS was performed with 1 μg/mL of mouse IgG1 versions of V312, TD1 or isotype control followed by 1:500 APC-F(ab’)2 goat-anti-mouse IgG (Jackson Immunoresearch #115-136-071). V312 and TD1 only gave a signal on MS4A4A-transfected cells (Supplemental Figure 3B). All antibodies were specific for their target.

### CRISPR KO

RNPs were formed by complexing 1.2mL custom sgRNAs (IDT) with 0.5mL Cas9 enzyme (IDT cat# 1081061) at RT for 10 minutes. sgRNAs were MS4A4A: AACCATGCAAGGAATGGAAC and TATTCATTCCTAGACTACCT; MS4A6A: CAGACTATCCAGATCTTGTG and AAGTGGCCTGTTTGACAGAC; NTC: CGTTAATCGCGTATAATACG and CATATTGCGCGTATAGTCGC. RNPs were then nucleofected into 1.2-5X10^6^ monocytes isolated the prior day using the P3 Primary Cell 4D-Nucleofector® LV Kit (Lonza #V4LP-3520) on program CM137. Cells were thereafter cultured in macrophage differentiation media for a further five to seven days before use. Knockout was ≥90% and was confirmed using primers against MS4A4A (TTGGCAATGATAGACGTCGC and CATCAGAAGCAAGGAGTGGC) and MS4A6A (AGATTCATGTTGCTGCCACC and TCAGCCAAGAATTACCCTGG).

### PCR

RNA was extracted from cells using an RNEasy Plus Mini kit (Qiagen #74134). cDNA was made using a Superscript IV VILO MM kit (Invitrogen Cat# 11756050) on a Veriti thermocycler (Thermo Scientific). Quantification of transcript was performed using Taqman MM (Thermo-Fisher # 4444554) with primers to MS4A4A (Hs00254780-m1; FAM), MS4A6A (Hs01556747-m1; FAM) and GAPDH (Hs02758991_g1; VIC-PL) on Quant Studio 6 (Applied Biosystems).

### Immunofluorescence

4X10^4^ macrophages/well were plated on poly-D-lysine coated plates. After attaching, cells were washed then fixed with 4% PFA (Electron Microscopy Sciences #15710), then washed again. Cells were blocked for 1 hour at room temperature in blocking buffer (10% goat serum (MP Biomedicals #ICN19135680) + 22.52μg/mL glycine (Sigma #G8898) + PBS + 0.1% Tween-20), then incubated overnight at 4C in primary antibody diluted in antibody incubation buffer (10% goat serum + 0.1% tween-20 in PBS). Cells were washed, then incubated for 1 hour at room temperature in secondary antibody diluted in antibody incubation buffer. After washing, cells were stained with DAPI for 10 minutes, washed and then imaged on an Operetta microscope (Perkin-Elmer). Primary antibodies were mouse anti-MS4A4A (R&D #MAB7797SP; 1:250), rabbit anti-MS4A6A (Abcam #189983; 1:250). Secondary antibodies were Goat-anti-mouse-A488 (Invitrogen #A11001; 1:500) and goat anti-rabbit-A594 (Invitrogen #A11012; 1:500).

### sTREM2

A custom MSD was used to detect sTREM2. Multi-Array 96 well plates (Mesoscale Discovery Cat #L15XA-3) were coated overnight at 4C with 50 mL of 1 μg/mL anti-TREM2 (Alector; Clone 8F11) capture antibody. The next day, plates were washed three times in PBS+0.05% Tween-20, then blocked with 1% BSA (MP Biomedicals #820451) in PBS for two hours. After washing three times, 50 mL of sample diluted 1:2-1:10 depending on the experiment or human TREM2-Fc standard (R&D Systems #1828-T2-050) were added to the plate and incubated at room temperature for one hour with shaking. Plates were washed three times, then 50 mL of 100 ng/mL goat anti-human biotinylated TREM2 detection antibody (R&D Systems #BAF1828) was added. After shaking at room temperature for one hour, plates were washed and 50 mL SulfoTag Strepavidin (MSD #R32AD-1) diluted 1:2,500 was added to the plates. After shaking at room temperature for 30 minutes, plates were washed and 150 mL of 2X READ Buffer T (MSD #R92TC-1) was added. Plates were read on a Sector 600MM machine (MesoScale Discovery). Sample concentrations were interpolated from the standard using a 5PL logistic regression model.

### Viability assay

5X10^4^ cells were plated in duplicate and stimulated with serial dilutions of V312, TD1 or isotype control. Viability was measured after 48 hours with Cell Titer Glo (Promega #G7570) and luminescence was measured on an Envision 2105 luminometer (Perkin Elmer).

### Inflammatory soluble factor measurements

For in vitro studies macrophages or human microglia were stimulated in duplicate or triplicate for 48 hours with MS4A4A antibodies or isotype controls, then the supernatants were tested for CSF-1 (Mesoscale Discovery # K151XRK), OPN (R&D Systems #SOST00), IL1RA (Mesoscale Discovery # B21XP-3) or (Mesoscale Discovery # K151VLK). Human microglia were isolated from post-mortem brains of seven healthy and five AD brains (Charles River Laboratories). For in vivo studies on serum, kits for IL1RA, OPN and sCSF1R (R&D Systems #DY329) were first qualified before use. For in vivo studies on the CSF, Somascan was used.

### Metabolism

10^5^ macrophages were treated for 48 hours with MS4A4A antibodies or isotype controls in duplicates, then mitochondrial and glycolytic ATP generation was measured using a Glycolysis/Oxidative phosphorylation Assay kit (Dojindo #G270) according to manufacturer’s instructions.

### Lysosomal activity

To measure glucocerebrosidase activity, 10^5^ macrophages were treated for 48 hours with MS4A4A antibodies or isotype controls in duplicates, then cells were lysed in 60 mL N-per Neuronal Protein Extraction Reagent (Thermo-Fisher # 87792) + HALT protease/phosphatase inhibitor cocktail (Thermo-Fisher # 1861282). 100 mL supplemented McIlvaine buffer (McIlvaine buffer pH 4 + 3 mM sodium taurocholate (Sigma # T4009) and 25uL 50 μg/mL 4-Methylumbelliferyl β-D-glucopyranoside (Sigma #M3633) was added to 25uL lysate, then the plate was read over 60 minutes at 37C with kinetic monitoring at OD 355/460.

### Phagocytosis

Aggregated Ab was made using a Beta Amyloid (1-42), Aggregation Kit (rPeptide #A-1170-2). In brief, Ab was solubilized with 1% NH4OH then neutralized with 10X TBS pH 7.4 and incubated overnight at 37C. Aggregated Ab was next labeled with pHRODO Red Succinidimyl Ester (Thermo-Fisher #P36600) for 1 hour according to manufacturer’s instructions. Excess dye was removed with a methanol wash followed by four washes with HBSS. Line 01279 was treated for 48 hr with 1μg/mL TD1 or isotype control. Cells were then fed pHRODO-labeled Ab42 and imaged once per hour for 24 hrs on an Incucyte machine (Thermo-Fisher). After the final time points, cells were stained with Nuclear Green LCS1 (Abcam # ab138904) to quantify cell number. The total integrated intensity of the pHRODO signal was normalized to the counts of Nuclear Green, both assessed on Live cell Analysis software (Sartorius), to calculate Ab42 uptake over time.

### Proximity ligation assay

To detect the interaction between DAP12 and TREM2 interaction after MS4A4A antibody treatment, proximity ligation assay was performed following manufacturer’s instructions. Briefly, 5 × 10^4^ human macrophages were seeded in 96-well plate for overnight culture, following with a 48-hour treatment of 1 µg/mL MS4A4A antibody. After fixation with 4% paraformaldehyde, cells were incubated with primary antibodies against DAP12 (1:250 dilution, Santa Cruz Biotechnology #sc-133174) and TREM2 (1:250 dilution, Cell Signaling Technology #91068) overnight at 4oC. The interaction between DAP12 and TREM2 was detected by Biosciences Duolink in situ red starter kit (Sigma Aldrich). Cell images were captured with an Operetta High-Content Imaging System (Perkin Elmer), and high content analysis was performed through the Harmony 4.5 software.

### TREM2 signaling

Macrophages were primed with 1 µg/mL TD1 or isotype control for 48 hours, then cells were treated with an agonistic TREM2 antibody or isotype control for 5 minutes. Cells were lysed with cell lysis buffer (Cell Signaling Technology #9803) and endogenous levels of phosphorylated Syk at tyrosine 525/526 were detected with PathScan® Phospho-Syk (Tyr525/526) Sandwich ELISA Kit (Cell Signaling Technology #7970) following manufacturer’s instructions.

### NHP study designs

Two independent studies were conducted in cynomolgus monkeys (n=5/group) that were injected i.v. with either vehicle control or 80mg/kg of TD1, and serum, CSF and brain samples (approximately 1.0 - 0.5 cm3 each) from different brain regions (right frontal cortex and right hippocampus) were collected. For Study 1, animals were dosed on days 1 and 29 and takedown occurred on day 31. Blood samples for serum PK, sTREM2 and OPN analysis were collected on Days 1 at timepoints pre-dose, 0.5, 2, 6, 12, 24, 48, 96, 168, 240, 336, 504, and 672 hours post dose, as well as Day 29 at timepoints 0.5, 2, 6, 12, 24, 48 hours post dose. CSF samples for PK and sTREM2. OPN, IL1RA analysis were collected pre-dose, 0.5, 2, 6, 12, 24, 48, 96 (Day 1 only), 168 (Day 1 only), 240 (Day 1 only), 336 (Day 1 only), 504 (Day 1 only), and 672 (Day 1 only) hours post dose. Lysates from a hemi-brain were tested for TREM2, OPN and CSF1R. RNA-seq was performed on microglia isolated from the other hemi-brain. For study 2, doses were administered on Days 1, 8, 22, and 50 and takedown occurred on Day 51. Brain samples were used for IHC analysis.

### Detection of MS4A4A antibodies

Anti-MS4A4A antibody in cynomolgus monkey CSF were measured using a custom Gyrolab assay based on a stepwise sandwich format with a biotin labeled Goat anti-human IgG (Cat no. NBP1-74983; Novus Biologicals) as the capturing reagent and an Alexa-fluor® 647 labeled anti-human IgG (Cat no. 2049-31; Southern Biotech) as the detection reagent. Standards, quality controls, and samples, were diluted in Rexxip HN buffer (Gyrolab #P0004996, Gyrolab). Anti-MS4A4A present in sample was captured by the biotinylated anti-human IgG bound to the streptavidin coated affinity column of the Gyrolab bioaffy CD (Gyrolab #P0004180 / P0004253). Captured Anti-MS4A4A antibody was detected by the fluorescently labelled anti-human IgG antibody. The intensity of the fluorescent signals produced was proportional to the amount of Anti-MS4A4A present in the samples. Data was analyzed using Gyrolab xP and Excel.

### Brain sample processing

Frozen brain samples were lysed with Neuronal Protein Extraction Reagent (N-Per) (Thermo Scientific # 87792) and Halt Protease inhibitor (Thermo Scientific # 1861278) on ice for 20 minutes according to manufacturer instructions. Samples were centrifuged and supernatants were transferred to a new tube and stored at −80°C until further analysis. The total protein concentration in each sample was measured by BCA protein analysis kit (Thermo Scientific # 23225,) according to manufacturer instructions. The protein concentration values were used to normalize analyte concentrations measured in brain tissues.

### Microglia isolation protocol

Fresh NHP tissue (approximately 2 cm^3^) from frontal cortex was dissociated as previously described^55^. Cells were stained with CD11B antibody to label myeloid cells and washed prior to isolating live (DAPI negative), CD11b^+^ cells using the BD ARIA FUSION FACS. Cells were collected directly into lysis buffer for RNA extraction. RNA was extracted using Qiagen Plus Pico kit and libraries prepared using Lexogen 3’FWD kit according to manufacturer’s recommendation for low input RNA samples. All samples were quality checked at each stage for RNA and library quality and quantity prior to sequencing. Live CD11b^+^ cells (40K-75K) were FACS-sorted from dissociated cortical tissue using the protocol described above for bulk microglia dissociation and collected directly into lysis buffer. RNA was extracted immediately and stored at −80°C for long term storage. An aliquot of RNA was tested for quality (typically RIN>7) and quantity in Agilent Bio Analyzer or Tape station format.

### RNA-seq data processing for NHP

Sequencing libraries were prepared as per the instructions for the low input protocol in Lexogen Quant Seq 3’ end FWD library prep. Libraries were checked for quality and quantity and sequenced using Next Seq Midi platform. Raw data have been processed with the internal Alector pipeline, that includes the following steps: reads trimming, alignment to reference genome, sequence and alignment quality control, and differential expression analysis. Trim Galore tool has been used to remove low-quality ends from reads (with quality score Phred <28) and reads shorter than 25 bp. STAR aligner (version 2.7.2b) has been used to align reads to the macaque genome (Macaca_fascicularis_5.0.93 assembly). Sequence and alignment quality have been analyzed with FastQC and MultiQC tools respectively. All samples were of good quality with enough mapped reads (>2.5 million) and no outliers were detected on the PCA plot. Raw gene expression values were converted to Reads Per Million (RPM) and Wilcoxon rank-sum test was used to evaluate the significance between the isotype control and Ab treated groups (Figure 5J).

### Somascan data processing

CSF was used to run Somascan proteomic assay (SomaLogic), and median normalized data (medNormSMP) was used. In this format, normalization is applied to samples (vs. to Calibrator and Buffer in medNormInt data). We used SomaDataIO R package (https://github.com/SomaLogic/SomaLogic-Data) to transform expression data from medNormSMP file, as well as to extract covariates of interest (SampleID, SampleType, NormScale_20, SlideID) and probes features (AptName, SeqID, SeqIdVersion, SomaID, TargetFullName, Target, UniProt, EntrezGeneID, EntrexGeneSymbol, Organism, Units, Type, Dilution, PlateScale.Reference, and Dilution2). Expression data was further log2 transformed, and linear model was used to evaluate differentially expressed proteins in treated and control samples.

### Mice for xenotransplantation

App^NL-G-F^ Rag2^−/−^ Il2rγ^−/−^ hCSF1^KI^ mice (strain C57BL/6J with BALB/c) ^4^ were used to study the effect of MS4A4A antibody *in vivo* in an AD relevant model. This knock-in model contains the humanized Aβ region, including three known AD pathogenic mutations (the Swedish “NL”, the Iberian “F”, and the the Arctic “G”)^40^. Mice accumulate Aβ plaques and show phenotypical changes (deficits in learning, memory and attention) from 6 months of age^59, 60^. Mice are also immunocompromised due to the lack of both Rag2 and Il2rγ and are permissive to the proliferation of human microglia cells due to the expression of the human colony-stimulating factor 1 (CSF1) gene sequence. Mice were housed at the University of Antwerp (Belgium) in a pathogen-free (SPF) facility in groups of 2-5 animals per cage, with access to food and water *ad libitum* and with a 14/10-hour light/dark cycle at 21 °C and 32% humidity. All experiments were conducted according to protocols approved by the Ethical Committee of Laboratory Animals of the University of Antwerp, Belgium (Ethische Commissie Dierproeven approval no. 2021-13) following local and European guidelines.

### Differentiation of microglial progenitors for transplantations

The human induced pluripotent stem cell (iPSC) line SIGi001-A (Sigma-Aldrich), was differentiated into microglial precursors and transplanted into the brain of newborn pups (P4) following the MIGRATE (microglia in vitro generation refined for advanced transplantation experiments) protocol^49^. Briefly, iPSC was plated and maintained in human matrigel-coated six-well plates and in mTeSR1 media until reaching ∼70–80% confluence. Once confluent, cell colonies were dissociated into single cells and plated into U-bottom 96-well plates at a density of ∼15,000/well in mTeSR1 medium with the addition of BMP4 (50 ng ml^−1^), VEGF (50 ng ml^−1^) and SCF (20 ng ml^−1^) for 4 days. On day 4, embryoid bodies were transferred into six-well plates (∼20 embryoid bodies per well) in complete X-VIVO medium (Lonza) supplemented with SCF (50 ng ml^−1^), M-CSF (50 ng ml^−1^), IL-3 (50 ng ml^−1^), FLT3 (50 ng ml^−1^) and TPO (5 ng ml^−1^) for 7 days with full medium change on day 8. On day 11, differentiation medium was replaced with complete X-VIVO (Lonza) complemented with FLT3 (50 ng ml^−1^), M-CSF (50 ng ml^−1^) and GM-CSF (25 ng ml^−1^). On day 18, human microglial precursors were collected and engrafted into the mouse brains (0.5 million cells per pup) as previously described^39^. Before transplantation (P4), endogenous mouse microglia depletion was achieved by 2 consecutive (P2 and P3) intraperitoneal injection of the CSF1R inhibitor BLZ945 (dissolved in 20% (2-hydroxypropyl)-β-cyclodextrin (Sigma-Aldrich) at a dose of 200mg kg^−1^ body weight^37, 39^.

### Xenotransplantation treatment regimen

Six months old xenotransplanted App^NL-G-F^ Rag2^−/−^ Il2rγ^−/−^ hCSF1^KI^ mice (males and females), received intraperitoneal administration of either V312 (n=10) or isotype control (n=8) every other week over a period of 3 months (for both, 6 doses of 80mg Kg^−1)^.

### Human microglia isolation from mouse brain for single-cell transcriptomics

Twenty-four hours after the final dose, mice were sacrificed with an overdose of sodium pentobarbital and immediately perfused with ice-cold 1 × DPBS (Gibco) supplemented with 5 U of heparin (LEO). After perfusion, one hemisphere of each mouse brain without cerebellum and olfactory bulbs was placed in fluorescence-activated cell sorting (FACS) buffer (1x PBS, 2% FCS and 2 mM EDTA) with 5 μM actinomycinD (ActD; Sigma) and processed for scRNA sequencing as previously described^48^. Briefly, brains were mechanically and enzymatically dissociated using Miltenyi Neural Tissue Dissociation Kit (Miltenyi) supplemented with 5 μM ActD. Next, the samples were filtered through a 70-μm strainer (BD2 Falcon), washed in 10 ml of ice-cold FACS buffer with 5 μM ActD and spun at 300*g* for 15 min at 4 °C. ActD was removed from the myelin removal step to prevent toxicity derived from long-term exposure^48^. Following the dissociation, myelin was removed by resuspending the pelleted cells in 30% isotonic Percoll (GEHealthcare) and centrifuging at 300*g* for 15 min at 4 °C. Accumulating layers of myelin and cellular debris were discarded, and Fc receptors were blocked in FcR blocking solution (1:10, Miltenyi) in cold FACS buffer for 10 min at 4 °C. Cells were then washed in 5 ml of FACS buffer and pelleted down. Samples were hashed individually using Total-Seq A^TM^ cell hashing antibodies (1:500, BioLegend) and incubated along with the following antibodies: PE-Pan-CD11b (1:100, Miltenyi), BV421-mCD45 (1:100, BD Biosciences), APC-hCD45 (1:50, BD Biosciences), and viability dye (1:2,000, eFluor780, Thermo Fisher Scientific), in cold FACS buffer for 30 min at 4 °C. After incubation, cells were washed, and the pellet was resuspended in 500 μl of FACS buffer and passed through a 35-μm strainer before sorting. For sorting, the cell suspension was loaded into the input chamber of a MACSQuant Tyto HS Cartridge and human cells were sorted based on CD11b and hCD45 expression at 4 °C. FACS data were analyzed using FCS Express 7 software.

### Single-cell library preparation and sequencing

Between 20,000–100,000 human microglia (CD11b+, hCD45+) from each mouse brain were sorted on the MACSQuant Tyto and diluted to a final concentration of 2,000 cells per μl for single-cell RNA sequencing. A total of 3300 human microglia cells per mouse brain were pooled and loaded onto the Chromium Next GEM Chip G (PN 1000120). The DNA library preparations were generated following the manufacturer’s instructions (CG000315 Chromium Next GEM Single Cell 3′ Reagent Kits v3.1). In parallel, the hashtag oligo libraries were prepared according to the manufacturer’s instructions (BioLegend, Total-Seq A Antibodies and Cell Hashing with 10x Single Cell 3′ Reagent Kit v3 3.1 Protocol) using 15 cycles for the index PCR. Four libraries for a total of 18 samples were sequenced targeting a 90% messenger RNA and 10% hashtag oligo library (45,000 reads per cell), on a MGIseq-2000 (Illumina) platform with the recommended read lengths by the 10X Genomics workflow.

### Analysis of single-cell RNA sequencing datasets

The FASTQ files were aligned by Cellranger (v.7.1.0) against the human genome database (GRCh38, Ensembl 98, refdata-gex-GRCh38-2020-A). Raw count matrices were imported in R (v.4.3.2) for data analysis. The datasets were analyzed using the Seurat R package pipeline (v.4.3.0.1). Cells of multiple mice were pooled in each library (Supplemental Table 1). Demultiplexing of single cells was executed using the MultiseqDemux()^7^ function, with the parameters autoThresh set to TRUE and maxiter set to 20. Mice MG4 and MG21 were excluded from further analysis, because almost no sequencing reads were generated for their hashes, hence not allowing an accurate assignment of their cells. After demultiplexing, it was observed in library 31 that mouse MG221 contributed approximately 6,776 cells, while the remaining mice contributed on average 980 cells. To avoid analysis bias, we down sampled the number of cells of mouse and 980 cells were randomly selected from MG221 and retained for further analysis. Subsequently, low-quality cells were removed from each library, which were identified as having a high and/or low library size; a high percentage of reads mapping to mitochondrial genes; or cells with a low number of detected genes. These thresholds were determined separately for each library based on the median absolute deviation as implemented in the isOutlier () function of the scuttle 1.12.0 package^61^. After quality control, each library was individually normalized using NormalizeData (). For all libraries, the 2,000 most variable features were selected with FindVariableFeatures () for downstream integration. A list of common integration anchors across libraries was selected with FindIntegrationAnchors () that was subsequently used for integration. Finally, integration was performed with IntegrateData () to correct for any potential library batch effect. This resulted in a dataset of 23,120 genes and 13,057 single cells divided over 16 mice.

A first analysis was performed to identify and retain only the microglia. This was done by performing a standard clustering analysis in Seurat on the integrated dataset, i.e. ScaleData (), RunPCA (), FindNeighbours (), FindClusters with resolution parameter set to 0.5, and finally RunUMAP (). This analysis resulted in 12 clusters, of which four were removed: a cluster with expression of *MRC1* and *CD163* (macrophages); one with expression of *CDK4*, *PCNA*, *TK1* (cycling cells); one with expression of *FLT1* (endothelial); and one with expression of *PLP1* (oligo). Cells belonging to these four clusters were removed and the data was integrated in the same manner as described previously resulting in an integrated dataset consisting of only microglia cells.

A second analysis was performed on this integrated dataset. The genes in the integrated object were scaled with ScaleData () and subsequently RunPCA () was run. An appropriate number of principal components (15 in total) was selected based on an elbow plot, capturing the most relevant sources of variation while avoiding excessive noise. After clustering, the mouse MG223 was considered an outlier and removed, because 40% of its cells fell into the IRM cluster (*ISG15*, *IFI6*, *IFITM3*, *IFIT1*, *IFIT2*, *IFIT3*). Next, integration steps were performed once more as described previously. This led to the final dataset which consisted of 11,167 high quality cells divided over 15 mice.

This integrated data set was processed as previously. Different resolutions for the clustering were tested, and the final clustering was performed with a resolution of 0.6. This initially led to the 8 clusters of which some were merged afterwards. The resulting 6 clusters corresponding to the transcriptional states human microglia we previously reported in xenotransplanted micrgolia in response to Aβ pathology^48^. The clusters were annotated based on their expression of the top ten markers, as previously identified^48^. Additionally, module scores were inspected using the AddModuleScore () function, considering features from the top 100 most differentially expressed genes in ^48^. These genes were selected based on fold change when comparing a given cluster to all other clusters. Lastly, the marker genes of each cluster were calculated with the wilcoxaux () function of the presto 1.0.0 package^62^. This led to the following annotation: HM (cluster 0, 5 and 7), HLA (cluster 1), CRM (cluster 2), DAM (cluster 3), IRM (cluster 4), and RM (cluster 6) (Supplementary Figure 8).

After obtaining this final clustering of the single cells at the cell state-level, we performed two analyses. First, a differential expression between the control and V312 treated mice was conducted with the FindAllMarkers () function with the logfc. threshold set to 0. A gene was called differentially expressed when the absolute value of the average log2FoldChange was larger than or equal to 0.2 and the adjusted p-value was lower than or equal to 0.05.

Our differentially expressed genes were compared to differentially expressed genes in the acute response from bulk RNA-seq from Cadiz et.al.^50^ (Figure 1, Panel D). Cadiz et.al. performed their analyses on mice, so the mouse genes had to be converted to their human counterparts. This conversion was performed DIOPT, because it combines multiple prediction tools^63^. The differentially expressed genes for ^50^ were used as input and genes with “Yes” for “Best Score” and “Best Score Reverse” were used to capture the maximum number of counterparts.

### Histology

After perfusion, the other brain hemisphere of each mouse was fixed in 4% paraformaldehyde (PFA) overnight at 4 °C. After 24 hours, PFA was discarded, and the brain tissues were washed and kept in 1X PBS at 4 °C until further processing. Fixed brain tissue was embedded in 4% Top Vision Low Melting Point Agarose (ThermoFisher). Agarose brain blocks were cut coronally in 40 µM thickness sections with a vibrating microtome (Leica). Each sample was collected under free-floating conditions and stored in cryoprotectant solution (40% PBS, 30% ethylene glycol, 30% glycerol) at −20 °C. For the staining, sections were washed with 1X PBS and permeabilized for 15 min at room temperature in PBS with 0.1% Tween 20 (PBST). After permeabilization, sections were stained with X-34 staining solution (1 μM X34, Sigma-Aldrich), 20 mM NaOH (Sigma-Aldrich) and 40% ethanol for 20 min at room temperature. Sections were washed 3 times for 2 minutes with 40% ethanol and 2 times for 5 minutes with PBST. Sections were then blocked with 5% normal donkey serum in PBST for 1 hour at room temperature. To stain microglia, the primary and secondary antibodies in (Supplemental Table 2) were used. Four co-staining combinations were performed, all including X34 and hP2RY12 and one additional antibody between anti-CD9, anti-HLA, anti-LAMP2 and anti-Aβ (82E1). Primary antibodies were incubated for 16 hours at 4°C. Sections were washed with PBST then secondary antibodies were added and incubated for 2 h at room temperature. For the HLA staining, the signal was enhanced using a Tyramide Super Boost kit (ThermoFisher), slices were incubated for 5 minutes in tyramide working solution, the rest of the protocol was performed according to the manufacturer’s instructions. Stained sections were mounted with DAKO mounting medium (Agilent). Images (Z-stacks) at 20X magnification were taken with a Nikon Ti2 Automated Widefield microscope (Nikon). To measure the shift in microglial cell states at the site of Aβ plaques, we used a modified Sholl analysis where the fluorescent intensity of microglial markers hP2RY12, hCD9 and hHLA or of the phagocytic marker LAMP2 and of diffuse Aβ plaques was measured through concentric rings (annuli) of increasing diameter surrounding the X-34 plaque center. The analysis was performed using Fiji ImageJ 2.14.0/1.54f after determining a threshold for background correction. Intensities of each channel were scaled for comparison using *z*-score normalization of intensities proximally and distally from the plaque center, the means of the inner and outer 5-6 annuli were generated in GraphPad Prism v.10.2.1. The normalized means were checked for normality distribution using Shapiro-Wilk normality test in R and one- or two-tailed Welch’s t-test was performed in GraphPad in Prism v.10.2.1. Representative images were taken on a Nikon Ti2 Automated Spinning Disk microscope (Nikon) and for hCD9 on a LSM700 Confocal Laser Scanning Microscope (Zeiss).

---

**Supplemental Figure 1.**
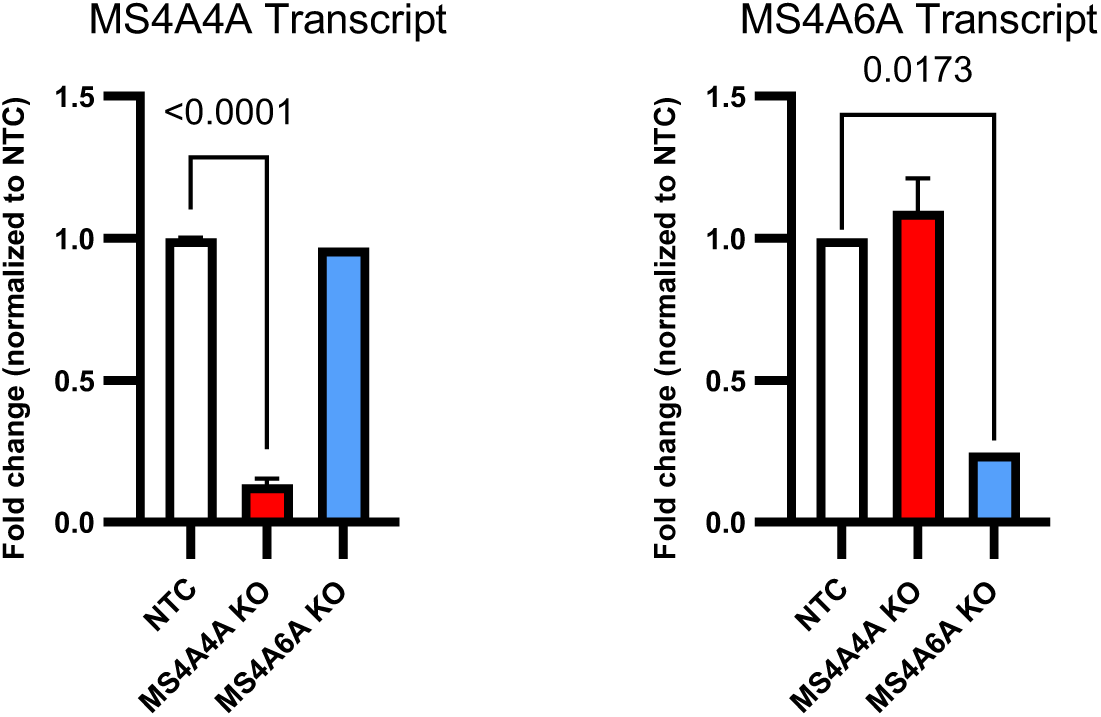
MS4A6A is regulated post-transcriptionally in MS4A4A KO macrophages. MS4A4A and MS4A6A transcript levels in NTC, MS4A4A KO and MS4A6A KO macrophages. Mean ± SEM of n=3 donors. p values were calculated with One-Way ANOVA.

**Supplemental Figure 2.**
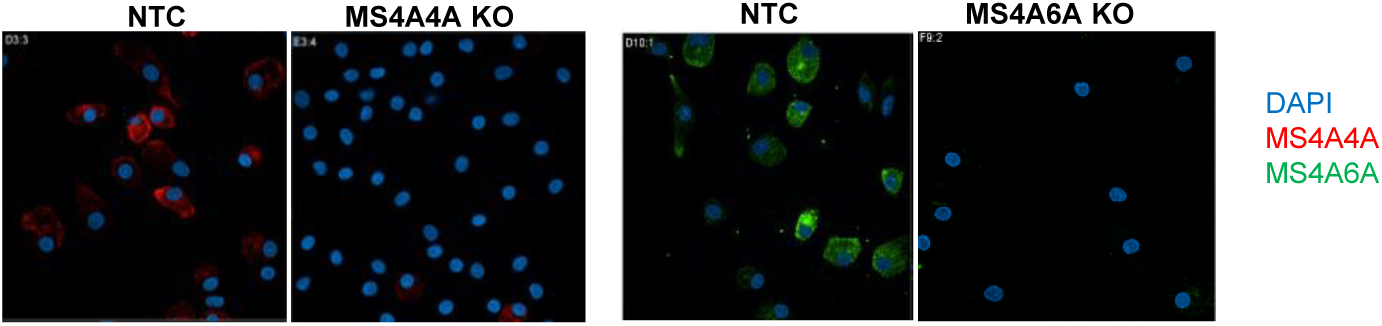
MS4A4A but not MS4A6A is localized on the cell surface. MS4A4A and MS4A6A were detected by immunofluorescence in NTC, MS4A4A KO and MS4A6A KO macrophages..

**Supplemental Figure 3.**
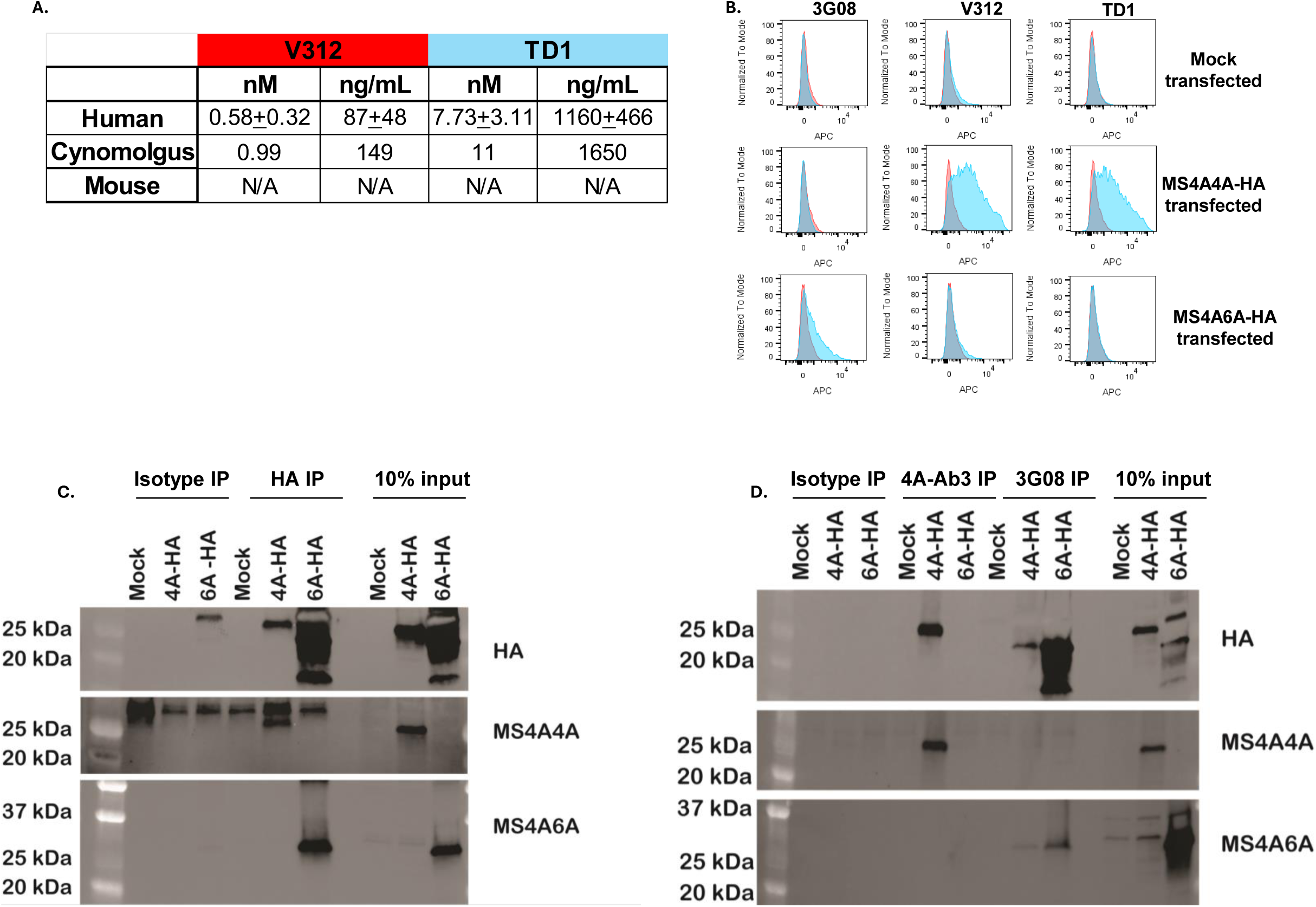
Characterization of MS4A4A- and MS4A6A-targeted antibodies used for treatment and reagents. (A) The apparent KD for binding of V312 and TD1 antibodies to U937 cells over-expressing human, cynomolgus or mouse MS4A4A was calculated. Data shows mean±SEM is shown for human MS4A4A from four independent experiments. (B-D) HEK293 cells were mock transfected or transfected with MS4A4A-HA or MS4A6A-HA. (B) Specificity of V312 and TD1 were tested by FACS. 3G08 was used as a control for MS4A6A expression. (C) Specificity of Western Abs for MS4A4A and MS4A6A were tested after IP’ing HA; or (D) specificity of IP antibodies for MS4A4A and MS4A6A were tested by using them for IP followed by immunoblotting for HA.

**Supplemental Figure 4.**
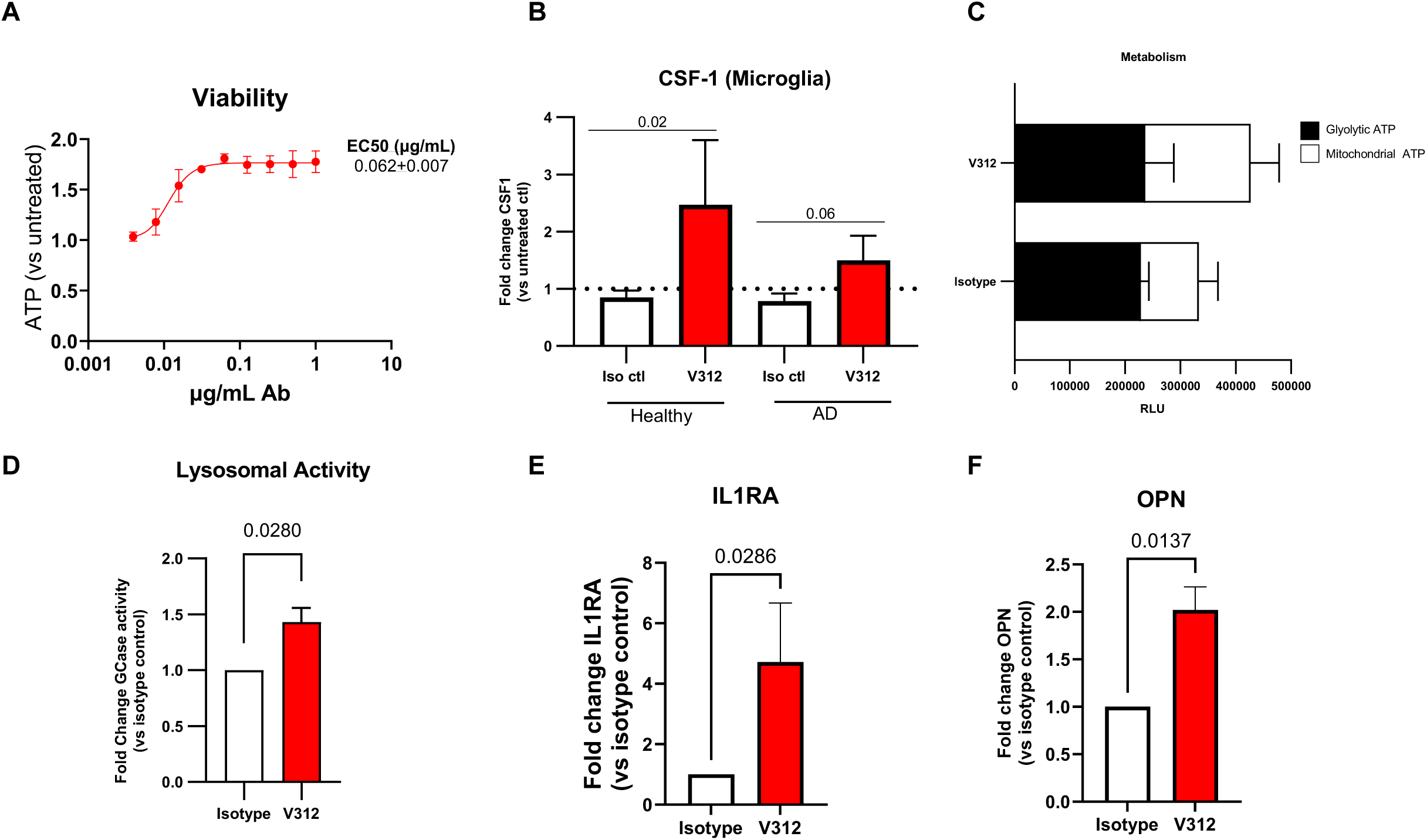
The MS4A4A-degrading antibody V312 regulates viability, metabolism, lysosomal function and cytokine production. (A) Viability (n=3 donors) was measured from macrophages after 48 hr treatment with V312 or isotype control, (B) CSF-1 production was measured after 48 hr treatment with 10 μg/mL V312 or isotype control from microglia isolated from postmortem brains of healthy controls (n=7) or AD patients (n=5). (C) glycolysis and oxidative phosphorylation (n=2 donors) was measured from macrophages after 48 hr treatment with 1 μg/mL V312 or isotype control. (C) was measured in macrophages (n-2 donors) after 48 hr treatment with 1 μg/mL V312 and TD1 or isotype control. (E) Line 01279 iPSC microglia were treated with with 1 μg/mL TD1 or isotype control. (D) Lysosomal activity was measured from macrophages (n=3 donors) 48 hrs after treatment with 10 μg/mL V312 and TD1 or isotype control (G) IL1RA (n=4 donors) and OPN (n=3 donors) was measured from macrophages after treatment for 48hrs with 1 μg/mL V312 or isotype control;. (A-F) Data shows mean±SEM. (A) EC50s were calculated with a 4PL non-linear regression. (B,D,E,F) p values were calculated with T tests.

**Supplemental Figure 5.**
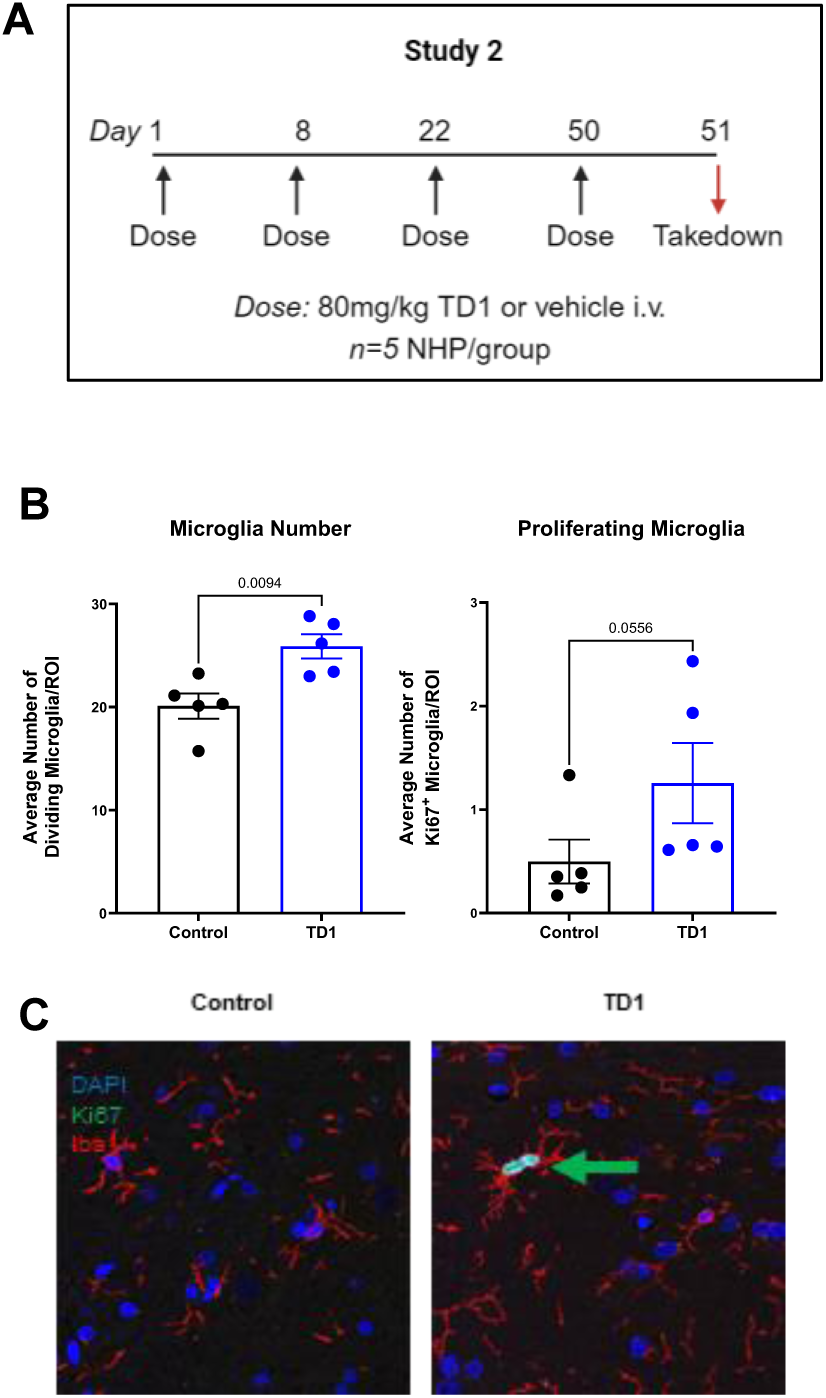
The MS4A4A-degrading antibody TD1 promotes microglia proliferation in non-human primate. (A) Design of NHP Study 2. (B) Proliferation index measured by Ki67 staining and total number of microglia 48hrs after final dose of TD1. (C) Immunoflurescence staing of Ki67 and Iba1. Data shows mean± SEM for n=5 animals/group. P values were calculated with T-test.

**Supplemental Figure 6:**
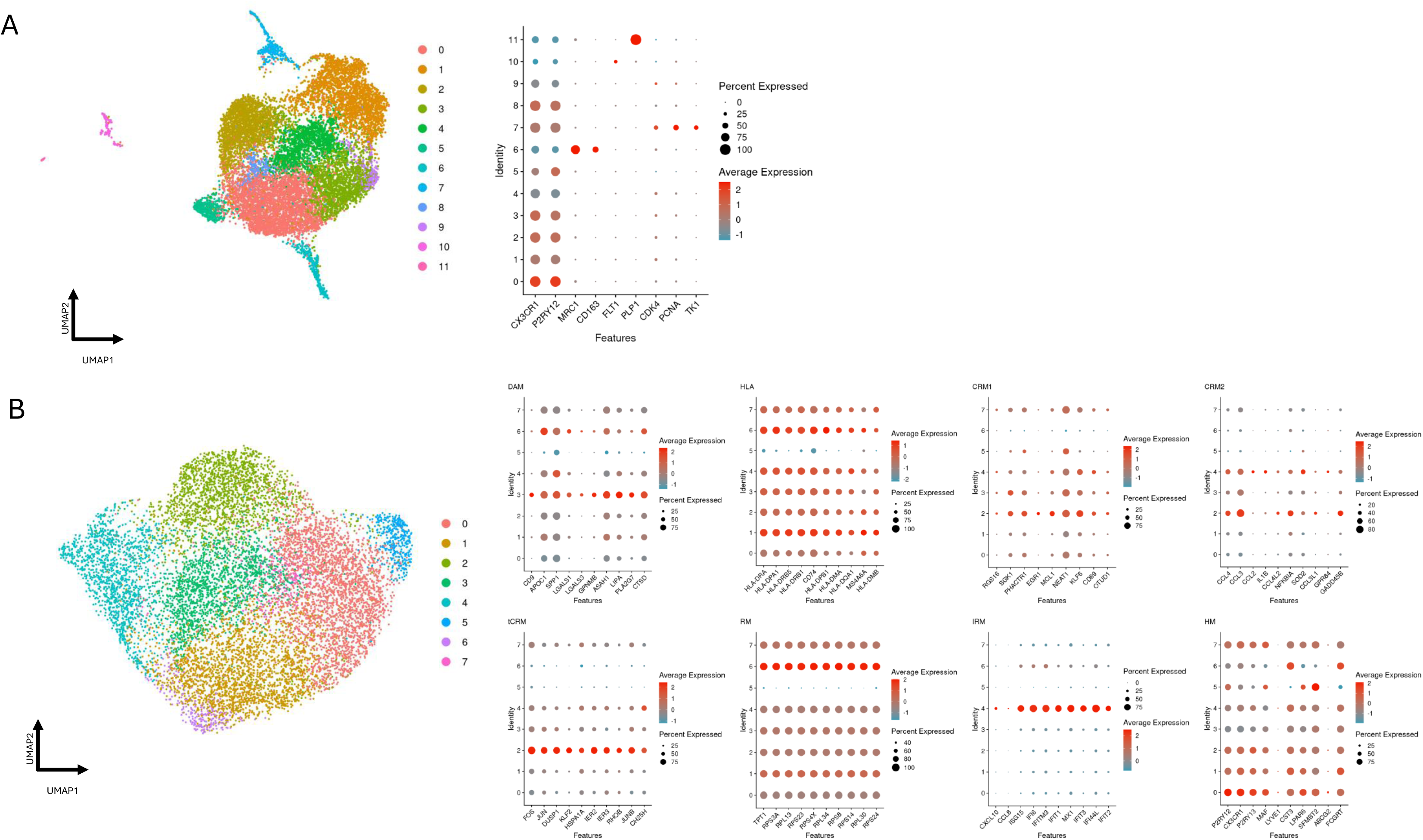
scRNA-Seq data processing workflow to identify microglia clusters. (A) UMAP of all 13,057 high quality cells after integration. Based on the dotplot, clusters 6 (macrophages), 7 (cycling), 10 (endothelial) and 11 (oligo) were removed. (B) UMAP of 11,167 high quality microglia cells after integration. The expression of the top 10 markers as in ^41^ are visualized with dotplots. This led to the following annotation: HM (cluster 0, 5 and 7), HLA (cluster 1), CRM (cluster 2), DAM (cluster 3), IRM (cluster 4), and RM (cluster 6).

**Supplemental Figure 7:**
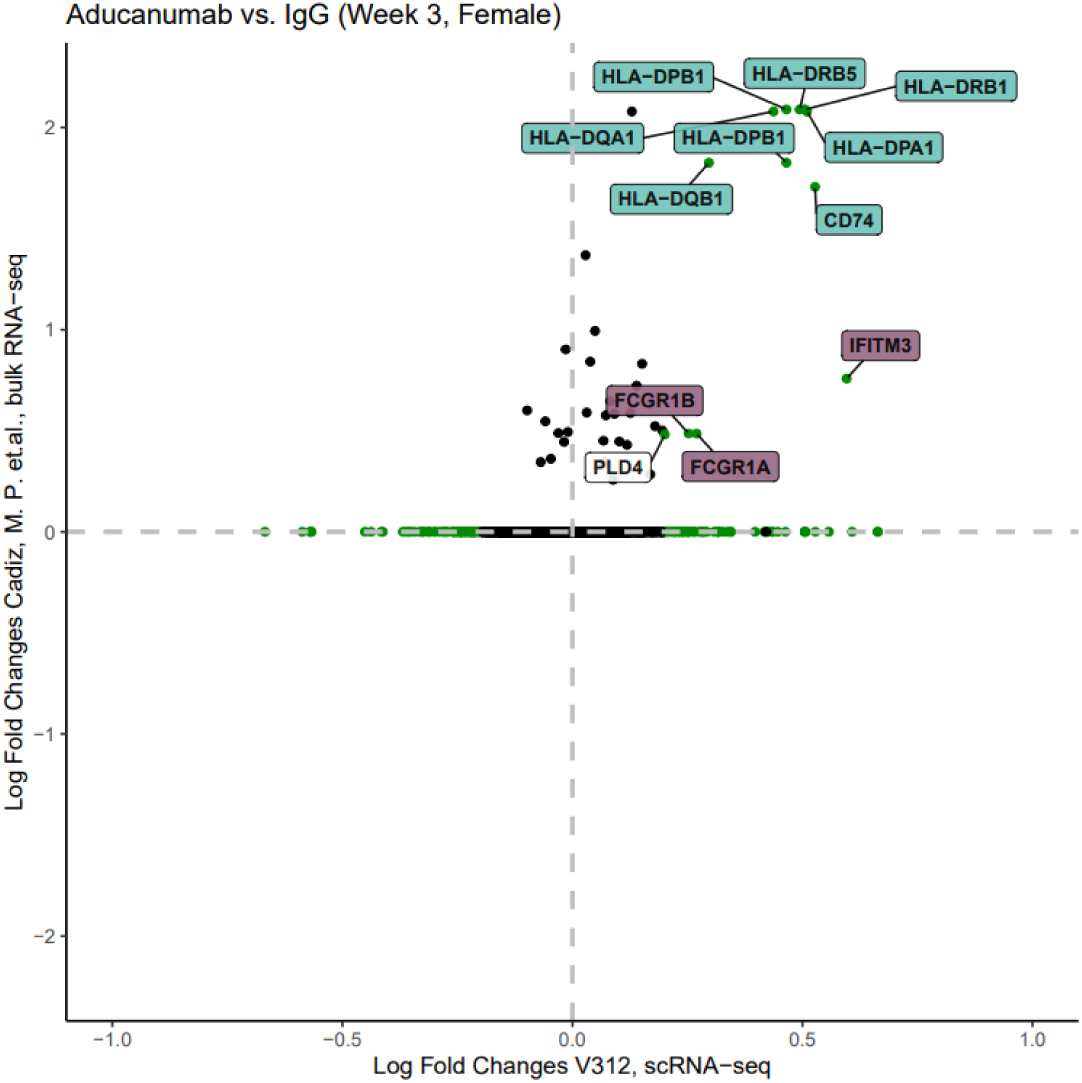
The MS4A4A-degrading antibody V312 can induce similar changes as an anti-Abeta antibody: Comparison of differentially expressed genes from V312-treated microglia with genes from aducanumab-treated animals in ^42^ showing enhanced MHC (HLA) response in female mice.

**Supplemental Figure 8:**
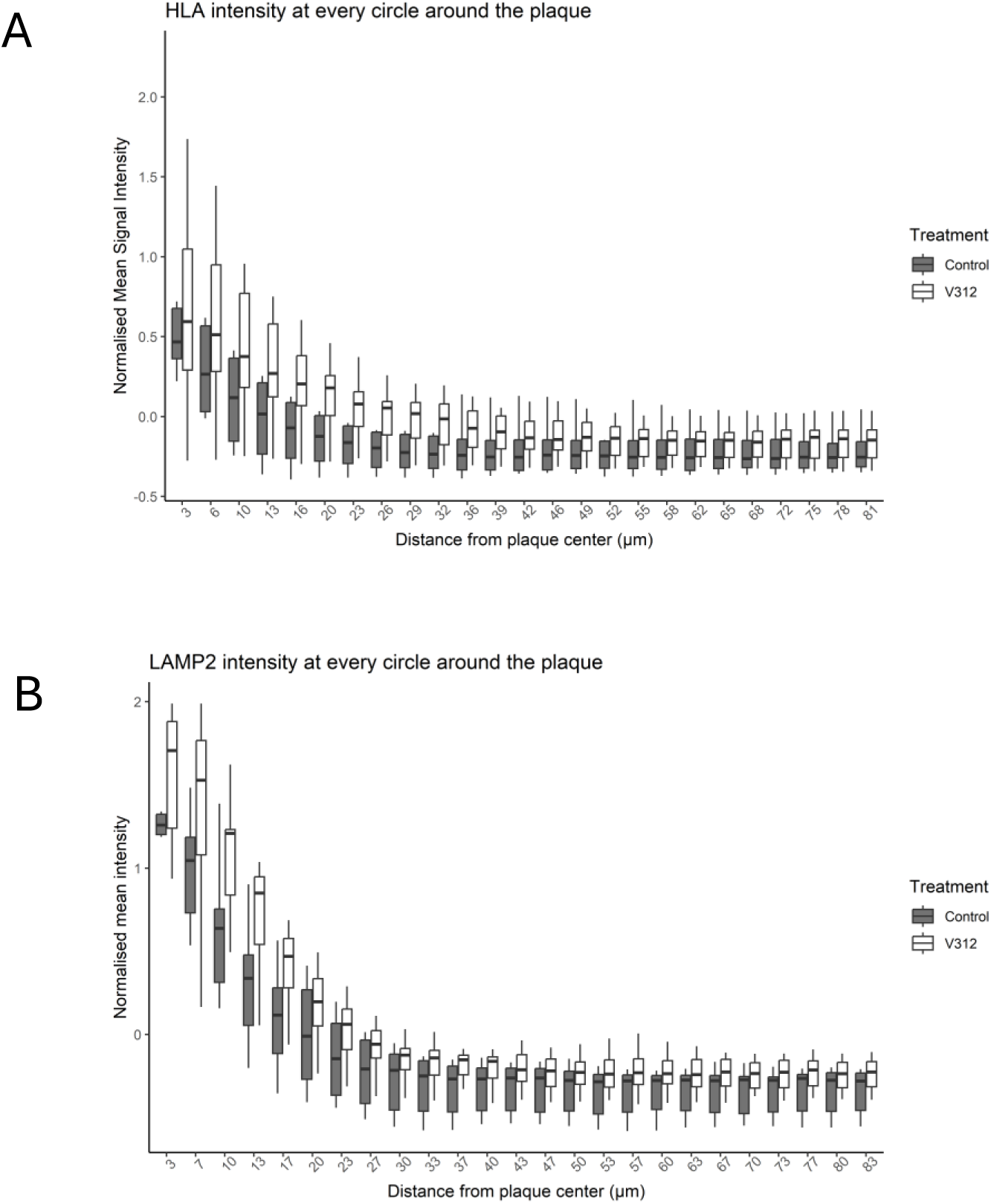
The MS4A4A-degrading antibody V312 alters microglia signature and activity regardless of distance to plaque: Fluorescence intensity distribution of (A) HLA and (B) LAMP2 over distance from plaques at each annulus. n=8 isotype control-treated mice and n=10 V312-treated mice

**Supplemental Figure 9:**
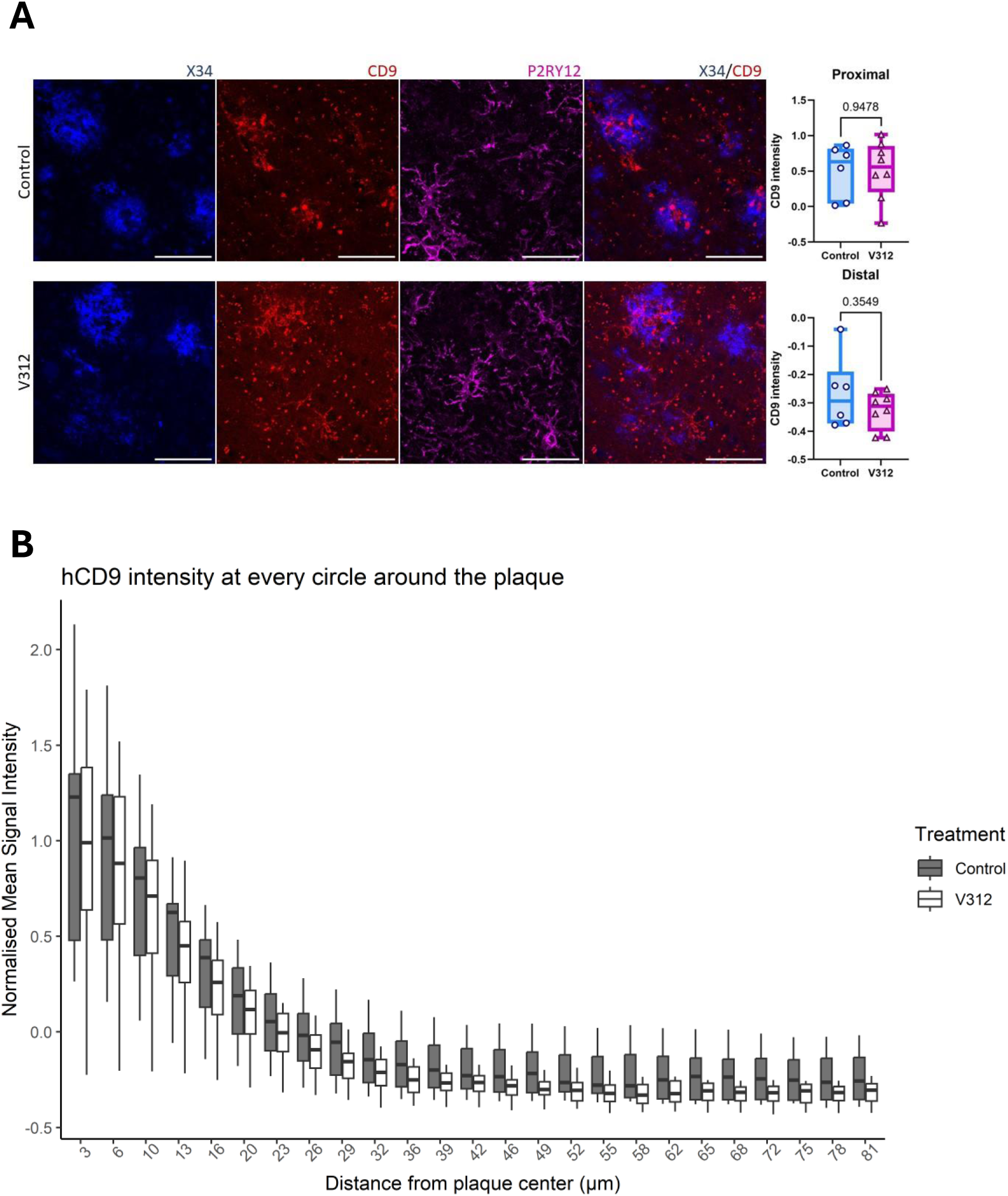
The MS4A4A-degrading antibody V312 did not alter the DAM microglia cluster at any distance from the plaques. (A) Representative images (scale bar 50 μm) and quantification of human microglia engrafted in the brain of App^NL-G-F^ Rag2^−/−^ Il2rγ^−/−^ hCSF1^KI^ mice after treatment with V312 or isotype control. Human microglia was labelled with human-specific antibodies for P2RY12 (HM microglia), and X-34 for Aβ plaques and hCD9 (DAM microglia); (B) Fluorescence intensity distribution of hCD9 over distance from plaques at each annulus. n=8 isotype control-treated mice and n=10 V312-treated mice.

**Supplementary Table 1:** ScRNA sequencing libraries specifications.

| LIB ID | Low Library Size | High Library Size | High Percentages Mitochondrial | Low Number Detected Genes |
| --- | --- | --- | --- | --- |
| LIB15 | 1818 | 27073 | 10 | 1278 |
| LIB16 | 1170 | 32724 | 15 | 880 |
| LIB31 | 1245 | 16367 | 4 | 856 |
| LIB46 | 1075 | 17192 | 6 | 788 |

**Supplementary Table 2:**
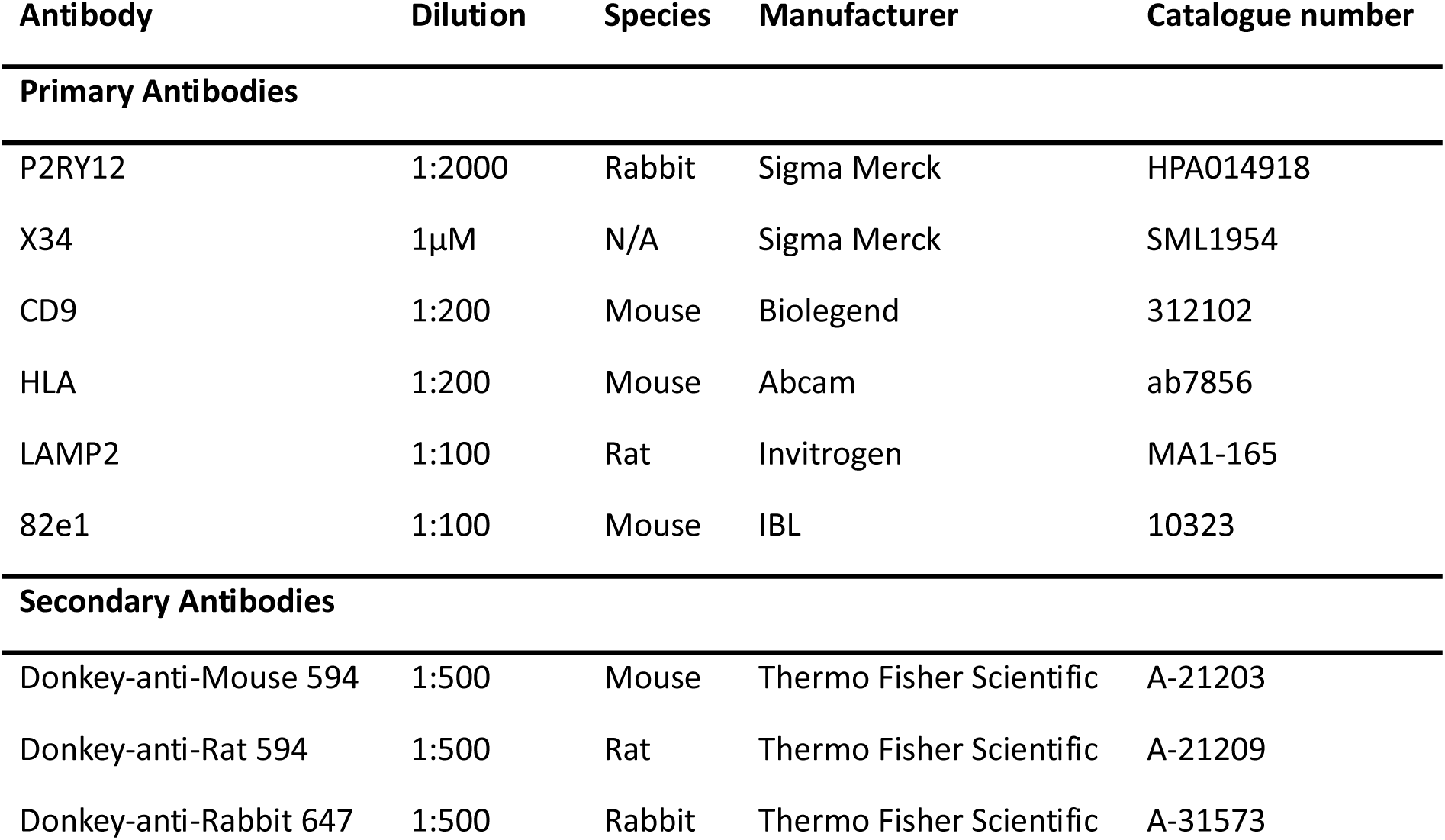
Antibodies used in immunohistochemistry.

